# A three-sodium-to-glycine stoichiometry shapes the structural relationships of ATB^0,+^ with GlyT2 and GlyT1 in the SLC6 family

**DOI:** 10.1101/2021.06.01.446649

**Authors:** Bastien Le Guellec, France Rousseau, Marion Bied, Stéphane Supplisson

**Affiliations:** Institut de Biologie de l’ENS (IBENS), Inserm, CNRS, École normale supérieure, PSL Research University, Paris, France

**Author notes:** **For correspondence:** (SS). These authors contributed equally to this work (Bastien Le Guellec, France Rousseau).

## Abstract

ATB^0,+^ (*SLC6A14*) absorbs all neutral and cationic amino acids in the distal colon and lung epithelia, and is part of the amino acid transporter branch I of the SLC6 family with GlyT1 (*SLC6A9*) and GlyT2 (*SLC6A5*), two glycine-specific transporters coupled to 2:1 and 3:1 Na^+^:Cl^−^, respectively. However, ATB^0,+^ stoichiometry that specifies its driving force and electrogenicity remains unsettled. Using the reversal potential slope method, here we demonstrate that ATB^0,+^-mediated glycine transport is coupled to 3 Na^+^ and 1 Cl^−^ and has a charge coupling of 2.1 *e*/glycine. ATB^0,+^ behaves as a unidirectional transporter with limited efflux and exchange capabilities. Analysis and computational modeling of the pre-steady-state charge movement reveal higher sodium affinity of the apo-ATB^0,+^, and a locking trap preventing Na^+^ loss at depolarized potentials. A 3 Na^+^/ 1 Cl^−^ stoichiometry substantiates ATB^0,+^ concentrative-uptake and trophic role in cancers and rationalizes its structural proximity with GlyT2 despite their divergent substrate specificity.

## Introduction

Several Na^+^-coupled amino acid transporters mediate the active uptake of glycine, which, despite its chemical simplicity - a nonessential amino-acid, without a side-chain or stereoisomer, and a membrane-impermeable zwitterion at physiological pH (***Chakrabarti, 1994***) - is nevertheless (1) a complex signaling molecule in the CNS as a fast inhibitory neurotransmitter and an agonist of the NR1 and NR3 subunits of the NMDA receptors (***Legendre, 2001***; ***Zeilhofer et al., 2012***; ***Johnson and Ascher, 1987***; ***Chatterton et al., 2002***; ***Grand et al., 2018***), (2) a key organic osmolyte for the regulation of cell volume in early embryos (***Dawson et al., 1998***), and (3) an essential metabolite for cell growth (***Wang et al., 2013***). Profiling studies have identified an increase in glycine one-carbon metabolism as an early warning signal of the rapid cell proliferation in cancers (***Jain et al., 2012***; ***Locasale, 2013***; ***Amelio et al., 2014***).

Three of these glycine transporters (GlyT1, GlyT2, and ATB^0,+^) and the proline transporter (PROT) are clustered in the amino-acid transporters branch (I) of the Solute Carrier 6 (SLC6) family, also named Neurotransmitter:Sodium Symporters (NSS) (***Bröer and Gether, 2012***). GlyT1 and GlyT2 correspond to the classic, high-affinity and glycine-specific “system Gly”, while ATB^0,+^ lacks substrate specificity (***Christensen, 1990***) and transports a broad variety of *α* and *β* amino acids, D- and L-amino acids, amino acid derivatives, and pro-drugs (***Sloan and Mager, 1999***; ***Winkle et al., 1985***; ***Hatanaka et al., 2001***; ***Nakanishi et al., 2001***; ***Hatanaka et al., 2002***, ***2004***; ***Anderson et al., 2008***). This broad specificity suggests that ATB^0,+^ has a distinctive substrate site from the one of GlyT1 and GlyT2, but nevertheless, we refer to ATB^0,+^ as a glycine transporter for simplicity.

GlyT1 and GlyT2 recapture and recycle glycine released at fast inhibitory synapses in the spinal cord, brainstem and cerebellum (***Legendre, 2001***; ***Eulenburg et al., 2005***; ***Zeilhofer et al., 2012***; ***Ankri et al., 2015***). GlyT2 is a specific marker of glycinergic neurons and operates like a unidirectional glycine pump as its 3 Na^+^/1 Cl^−^ stoichiometry provides an excessive driving force for influx (***Roux and Supplisson, 2000***; ***Supplisson and Roux, 2002***). GlyT2 inactivation disrupts the cytosolic accumulation and vesicular release of glycine (***Gomeza et al., 2003***; ***Rousseau et al., 2008***; ***Apostolides and Trussell, 2013***), and GlyT2^−/−^ knockout mice develop a severe hypoglycinergic syndrome (***Gomeza et al., 2003***) while mutations of human *SLC6A9* cause Hyperekplexia (OMIM:#614618), a rare neurological disorder with exaggerated startle reflexes (***Rees et al., 2006***; ***Carta et al., 2012***). In contrast, GlyT1 is as a bidirectional, 2 Na^+^/1 Cl^−^-coupled transporter that is expressed primarily on astrocytes and behaves as a buffer, sink, or source of extracellular glycine (***Zafra et al., 1995***; ***Supplisson and Bergman, 1997***; ***Huang and Bordey, 2004***; ***Aubrey et al., 2005***, ***2007***; ***Sipilä et al., 2014***; ***Shibasaki et al., 2017***). However, GlyT1 concentrative uptake is sufficient to specify the glycinergic phenotype of retinal amacrine cell (***Eulenburg et al., 2018***), and critical for the cell volume regulation of early mouse embryos (***Steeves et al., 2003***; ***Steeves and Baltz, 2005***). On the extracellular side, GlyT1 tunes the basal concentration and spillover of glycine (***Sipilä et al., 2014***; ***Ahmadi et al., 2003***) that gate NMDARs activation depending on their subunits composition, synaptic location, brain structure, and developmental stage (***Supplisson and Bergman, 1997***; ***Supplisson and Roux, 2002***; ***Tsai et al., 2004***; ***Martina et al., 2005***; ***Papouin et al., 2012***; ***Bail et al., 2015***; ***Ferreira et al., 2017***; ***Otsu et al., 2019***). GlyT1-deficit leads to an hyperglycinergic phenotype with hypotonia and motor disorders in GlyT1^−/−^-knockout mice and *Shocked* Zebrasfish-mutant (***Gomeza et al., 2003***; ***Tsai et al., 2004***; ***Cui et al., 2005***; ***Mongeon et al., 2008***; ***Hirata et al., 2010***; ***Eulenburg et al., 2010***). Mutations of human *SLC6A5* caused glycine encephalopathy with normal serum glycine (OMIM:#617301), a severe metabolic disease that produces hypotonia and respiratory failures in neonatal (***Kurolap et al., 2016***; ***Hauf et al., 2020***).

In comparison, the energetic and biophysical properties of ATB^0,+^ remain undercharacterized. ATB^0,+^ mediates electrogenic and concentrative uptakes of all neutral and cationic amino acids with Hill coefficients suggesting a 2 Na^+^ and 1 Cl^−^ stoichiometry (***Sloan and Mager, 1999***; ***Hatanaka et al., 2001***; ***Nakanishi et al., 2001***; ***Karunakaran et al., 2008***). ATB^0,+^ is expressed in the lung and distal colon epithelia, and in the mammary and pituitary glands (***Villalobos et al., 1997***; ***Sloan and Mager, 1999***; ***Ugawa et al., 2001***; ***Sloan et al., 2003***; ***Chen et al., 2020***). Mice lacking ATB^0,+^ are viable and with normal phenotype (***Babu et al., 2015***; ***Ahmadi et al., 2018***), but genome-wide association studies have identified SNPs in non coding regions of *SLC6A14* that are linked to obesity (***Suviolahti et al., 2003***; ***Corpeleijn et al., 2010***; ***Sivaprakasam et al., 2021***), male infertility (***Noveski et al., 2014***), and the phenotypic variation and severity of cystic fibrosis (***Sun et al., 2012***; ***Corvol et al., 2015***). In particular, the lack of ATB^0,+^ impairs intestinal fluid secretion in the context of cystic fibrosis and is responsible of Meconium Ileus at birth (***Ahmadi et al., 2018***; ***Ruffin et al., 2020***). Ganapthy’s group has demonstrated the implication of ATB^0,+^ in cancers, as one of the four amino-acid transporters with *SLC1A5*, *SLC7A5*, *SLC7A11* that are up-regulated in tumors for matching the increased demand in amino acids for cell growth, and more particularly for glutamine and glycine (***Bhutia et al., 2014***; ***Coothankandaswamy et al., 2016***; ***Sikder et al., 2017***; ***Sniegowski et al., 2021***).

Here, we investigated the stoichiometry and exchange mode of ATB^0,+^ in order to better evaluate its driving force and transport properties under physiological and pathological conditions. An additional motive was to understand the raison d’être of the surprising structural hierarchy between these three glycine transporters of the SLC6 family. We used the reversal potential of ATB^0,+^ in order to establish its stoichiometric coefficients by a thermodynamic method. Our results revise the apparent consensus for a 2 Na^+^ stoichiometry, and the charge/glycine and charge movement in glycine-free media confirmed the higher Na^+^ coupling. Finally, we show that 3 Na^+^-coupling reduces both the efflux and exchange of glycine in ATB^0,+^ and GlyT2-expressing oocytes, a property that is not shared by other 3 Na^+^-coupled transporters like the glutamate transporter EAAT2 who operates by a distinct elevator mechanism (***Kavanaugh et al., 1997***; ***Drew and Boudker, 2015***).

## Results

### Odd phylogenetic and functional relationships of ATB^0,+^ with GlyT1 and GlyT2

The position of ATB^0,+^ in the phylogenetic tree of SLC6 transporters (***Figure 1***A), closer to GlyT2 than this one is of GlyT1, seems at odds with their marked functional divergence in terms of specificity and affinity for glycine, as detailed below and in the next paragraph. Although the length of the common branch is short (0.14 for ATB^0,+^/GlyT2 *vs.* 0.07 for GlyT1/PROT, ***Figure 1***A), the same split is nonetheless observed for all Vertebrates orthologs despite a greater divergence in ATB^0,+^ sequences (***Figure 1***A-B). Indeed, the percentage of identities (ID) between orthologs is lower for ATB^0,+^ (39 % ID) than for GlyT2 (69.7 % IDs) and GlyT1 (60.3 % ID) as shown in ***Figure 1***B. A higher divergence is also detected in the more conserved transmembrane segments with 46.5 % *vs.* 77.3 %, and 70 % IDs, respectively ***Figure 1***–***Figure Supplement 1***A. Venn diagrams of private and shared residues confirm that mGlyT2 shares more IDs with mATB^0,+^ (n = 95) than with mGlyT1 (n = 76, ***Figure 1***C, ***Figure 1***–***Figure Supplement 2***) and the GlyT2/ATB^0,+^ pair shows a similar excess for each ortholog (+15.5±1.1 ID, n = 8, ***Figure 1***–***Figure Supplement 1***B). PROT was included in the analysis to count only for IDs specific of a glycine-transporter pair (GlyT2/ATB^0,+^: 55.4±1.1 ID, GlyT2/GlyT1: 39.9±1.5 ID, ATB^0,+^/GlyT1: 25.5±1.5 ID, n = 8, ***Figure 1***–***Figure Supplement 1***B).

**Figure 1.**
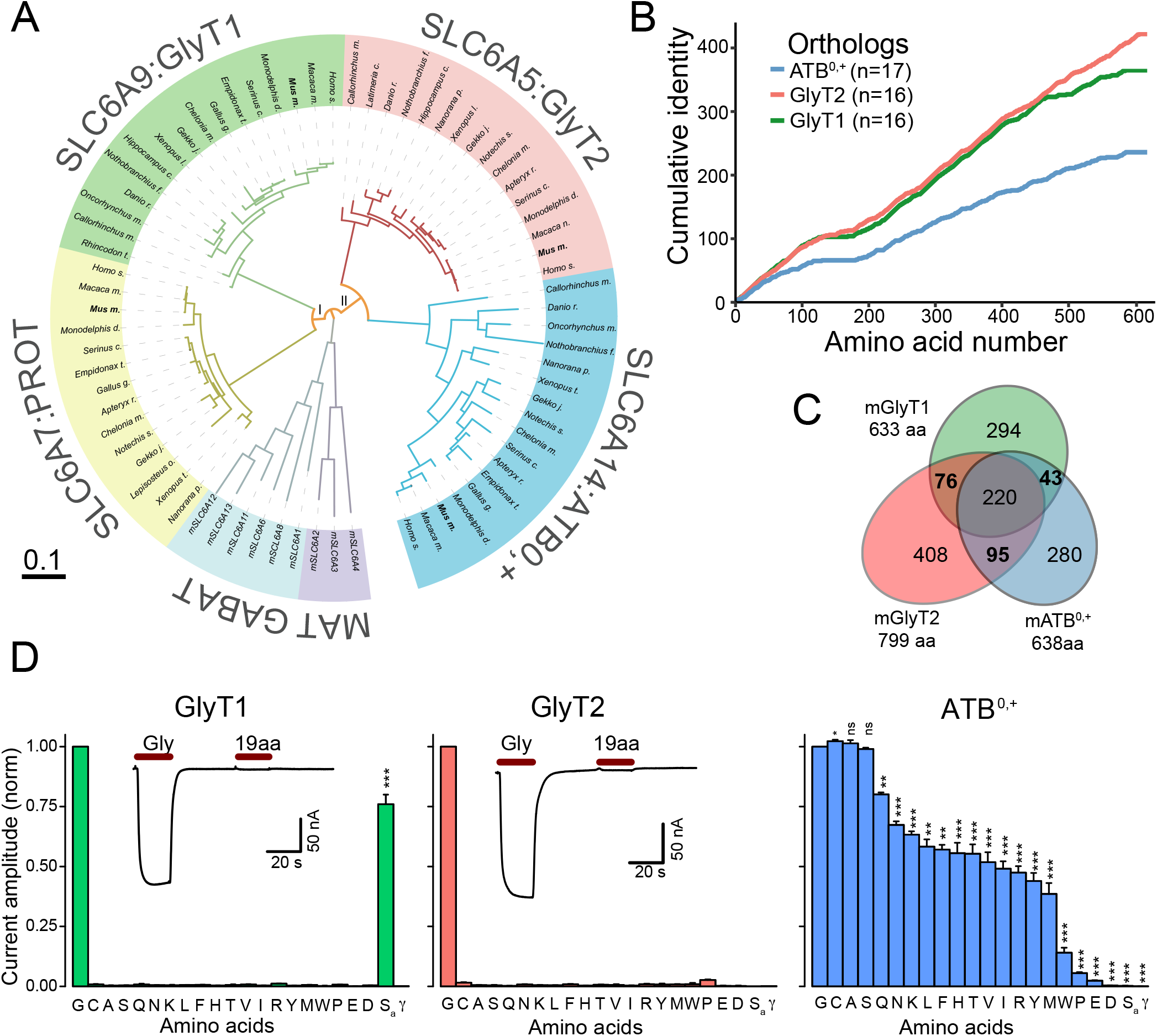
The divergent substrate specificity of GlyT1, GlyT2, and ATB^0,+^ does not predict their phylogenetic relationship in the SLC6 family. **(A)** The phylogenetic tree of amino acid transporter sequences of Vertebrates orthologs shows a split between two apparently unrelated pairs (I) PROT (n = 14) and GlyT1 (n = 16), and (II) GlyT2 (n = 16) and ATB^0,+^ (n = 17). The phylogeny includes the sequences of GABA transporters (GABAT: mGAT1 (*SLC6A1*), mTauT (*SLC6A6*), mCT1 (*SLC6A8*), mGAT3 (*SLC6A11*), mBGT1 (*SLC6A12*), and mGAT2 (*SLC6A13*), light blue) and monoamine transporters (MAT: mNET (*SLC6A2*), mDAT (*SLC6A3*), and mSERT (*SLC6A4*), purple) from *mus musculus* as outgroups. **(B)** The plot of cumulative identity for the same orthologs as in ***Figure 1***A shows that ATB^0,+^ (blue, n = 17) sequences are more divergent during Vertebrates evolution than GlyT1 (green, n = 16) and GlyT2 (red, n = 16). The cumulative count (+1 if all sequences shared the same residue) starts at the first shared-position in the alignment (R57 for ATB^0,+^, R309 for GlyT2 and R40 for GlyT1). **(C)** Area-proportional Venn diagrams of the number of shared and private residues for mGlyT1, mGlyT2 and mATB^0,+^ based on the sequences alignment shown in ***Figure 1***–***Figure Supplement 2***). GlyT2 shares more identity with ATB^0,+^ (n = 95) than with GlyT1 (n = 76) for all species examined ***Figure 1***–***Figure Supplement 1***). **(D)** Extreme divergence in substrate specificity. Individual bars represent the normalized current evoked by each amino acid in oocytes expressing GlyT1 (200 μM, n = 2-4, left panel), GlyT2 (200 μM, n = 4, middle panel) or ATB^0,+^ (1 mM, n = 6-7, right panel). Amino acids are identified by their single letter code, except sarcosine (S_a_) and GABA (*γ*). Insets show the absence of current when 19 amino acids (A,C,S,L,F,N,K,Q,H,V,T,Y,R,M,I,W,P,E,D at 200 μM are applied together compared to glycine (3.8 mM) in GlyT1-expressing oocytes (2.4 %, n = 6, p<0.001) and GlyT2-expressing oocytes (2.35 %, n=10, p=0.002). Error bars indicate SEM; paired t-test: p<0.01(***), p<0.01(**), p<0.05 (*). **Figure 1–Figure supplement 1.** Sequence identities among Vertebrates orthologs of ATB^0,+^, GlyT2 and GlyT1 **Figure 1–Figure supplement 2.** Sequences alignment of mATB^0,+^, mGlyT2 and mGlyT1

The substrate site of ATB^0,+^ is expected to differ substantially from the one of GlyT1 and GlyT2 as it can accommodate a large variety of amino acids size and side chains, with aliphatic, cationic, hydrophobic, or hydrophilic properties. Indeed, all the natural amino acids at the exception of glutamate and aspartate evoke an inward current in ATB^0,+^-expressing oocytes, in contrast to the absolute glycine specificity of GlyT1 and GlyT2 (***Figure 1***D). Therefore, other factors than a common substrate site for glycine must explain the structural proximity of ATB^0,+^ and GlyT2. Nevertheless, it is worth noticing that both transporters share the same intolerance for N-methyl-derivative while GlyT1 transports sarcosine (***Figure 1***D).

### ATB^0,+^ glycine-EC_50_ shows an anomalous voltage-dependency

Glycine evokes inward currents in ATB^0,+^-expressing oocytes that do not reverse up to +50 mV (***Figure 2***A), and are strictly dependent on Na^+^ and Cl^−^ (***Figure 2***–***Figure Supplement 1***). The current-voltage relationship is quasi-linear at saturating glycine concentration, indicating a weak voltage-dependency of its transport cycle similar to GlyT1 and GlyT2 (***Figure 2***B), with efold of 296, 313, and 254 mV, respectively.

**Figure 2.**
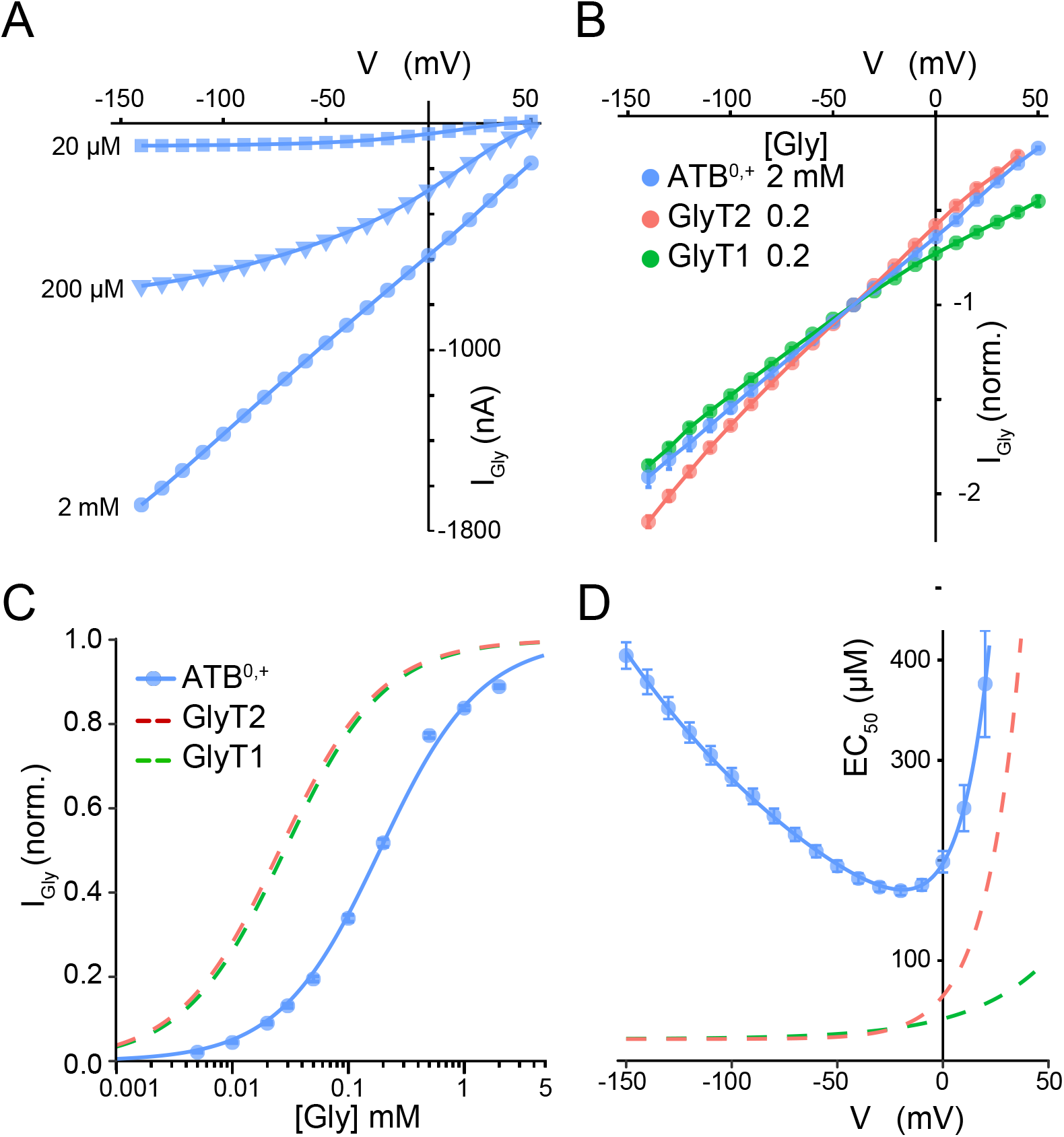
Weak and anomalous voltage-dependency of ATB^0,+^-mediated currents **(A)** Glycine evokes large inward currents in ATB^0,+^-expressing oocytes. Current-voltage (I-V) relationships of the glycine-evoked current (20 μM (square), 200 μM (triangle) and 2 mM (circle)). **(B)** Linear I-Vs recorded at saturating glycine concentrations for ATB^0,+^ (blue, [Gly]=2 mM, n = 10, slope = 0.92 % mV^−1^, R^2^ = 1.000, p<2 10^16^), GlyT2 (red, [Gly]=200 μM, n = 11, slope = 1.07 % mV^−1^, R^2^ = 0.998, p<2 10^−16^), and GlyT1 (green, [Gly]=200 μM, n = 11, slope = 0.87 % mV^−1^, R^2^ = 0.998, p = 1.78 10^13^). Currents are normalized by the absolute amplitude at −40 mV and linear regressions were performed on the −140–−40 mV voltage range. **(C)** Glycine dose-response curve for ATB^0,+^ (blue circles, V_*H*_ = −40 mV, n = 32). Individual curves were fitted by the equation: 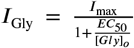, and then normalized by the maximal amplitude (*I*_max_). The solid line represents the fit of the normalized equation. The glycine EC_50_ of ATB^0,+^ (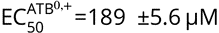, n=21) is higher than for GlyT1 (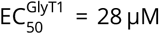, green dashed line) and GlyT2 (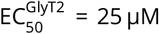, red dashed line) as previously determined in ***Roux and Supplisson*** (***2000***). **(D)** Anomalous voltage-dependency of 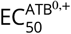 at hyperpolarized potentials (blue circles, n = 6). The solid line is the fit to the equation: 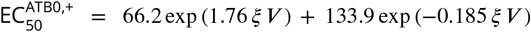. The dashed lines are fits for 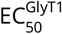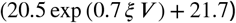 and 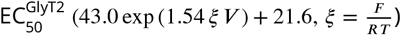 previously determined in ***Roux and Supplisson*** (***2000***). **Figure 2–Figure supplement 1.** ATB^0,+^ transport current is Na^+^ and Cl^−^-dependent

In contrast, ATB^0,+^ dose-response-curve for glycine shows a distinct and lower apparent affinity, with a more complex voltage-dependency than for GlyT1 and GlyT2 (***Figure 2***C-D) as previously reported in ***Roux and Supplisson*** (***2000***). In particular, ATB^0,+^-EC_50_ shows a minimum at −20 mV (171±4 μM, n = 6), and then increases at both, depolarized and hyperpolarized potentials, whereas GlyT2- and GlyT1-EC_50_ are voltage-independent in the negative range (***Figure 2***D, ***Roux and Supplisson*** (***2000***)).

On average, a saturating glycine concentration evokes ~ 6-fold larger currents in ATB^0,+^-expressing oocytes than for GlyT2 and GlyT1 (1450.0±98.8 pA (n = 65) *vs.* 250.0±29.7 pA (n = 48), and 245.2±17.3 pA (n = 85) at *V_H_* =−40 mV for 1–2 mM (ATB^0,+^) and 200 μM (GlyT2, GlyT1), respectively, ***Figure 3***A). Therefore, we examined whether the membrane-expression, charge per glycine, and turnover-rate can account for ATB^0,+^ larger electrogenicity.

**Figure 3.**
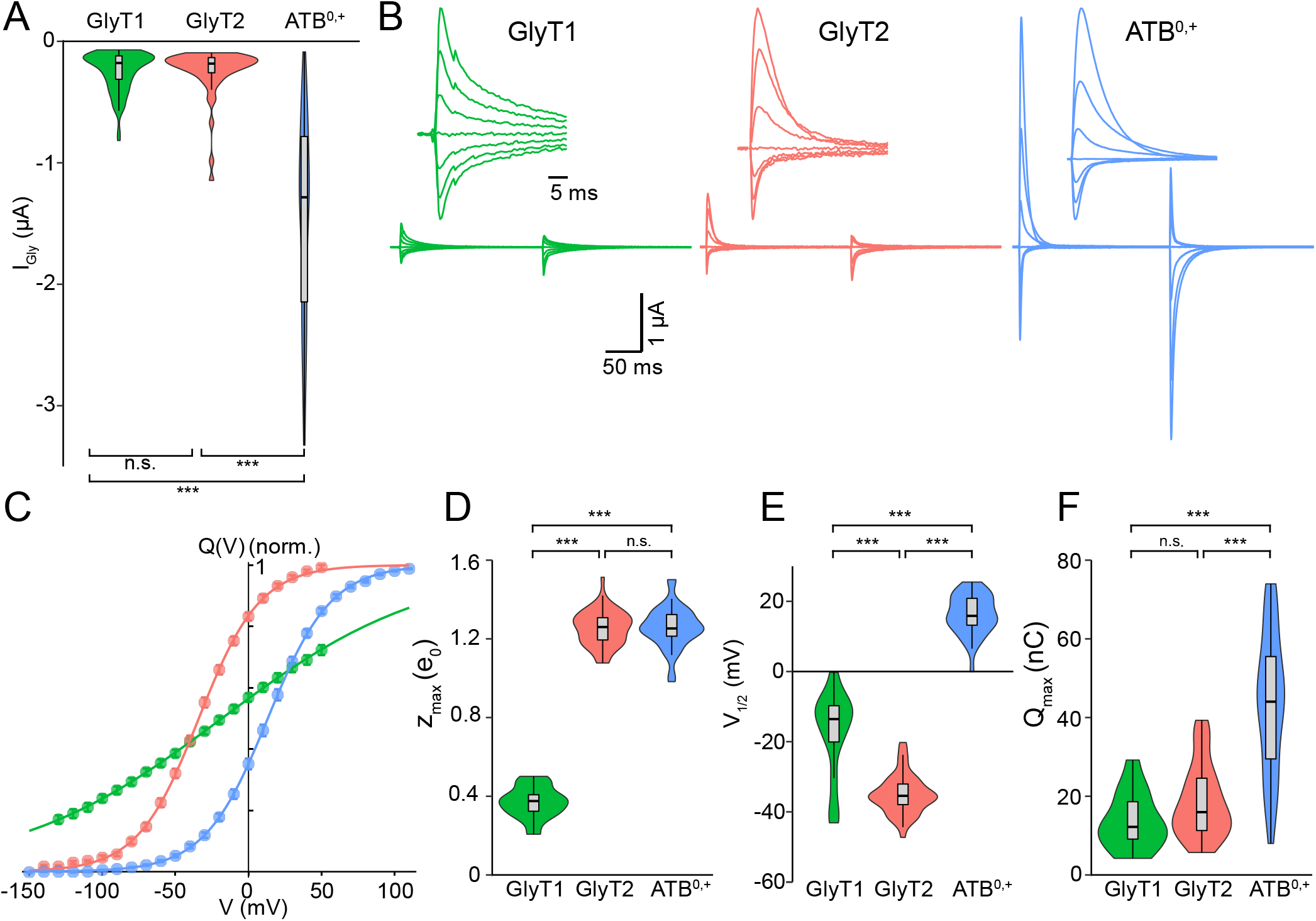
Presteady-state currents and the charge movement of ATB^0,+^ reveals a Na^+^-dependent convergence with GlyT2 **(A)** combined violin and box plots of transport current amplitudes recorded at −40 mV for different saturating glycine concentrations, that is 200 μM for GlyT1 (245.2±17.3 nA, n = 85) and GlyT2 (250.6±29.7 nA, n = 48) and 1–2 mM for ATB ^0,+^ (1450.0±98.8 nA, n = 65). **(B)** Representative presteady-state currents recorded in oocytes expressing GlyT1 (left, green), GlyT2 (middle, red) and ATB^0,+^ (right, blue) expressing oocytes. The range of voltage steps varied from +110 mV (ATB^0,+^) or +50 mV (GlyT1, GlyT2) to −150 mV by decrements of 10 mV. For clarity, only seven traces are shown, from +50 mV to −130 mV by decrements of 30 mV. Transporter currents were isolated by subtracting the traces recorded with the same voltage step protocol in the presence of specific inhibitors (10 μM ORG24598 (GlyT1); 5 μM ORG25543 (GlyT2); 1 mM *α*-MT (ATB^0,+^)). Any residual steady-state components have been subtracted for clarity, and insets are scaled to the maximal amplitude recorded at the onset of the voltage steps. **(C)** The charge movement is the time integral of the relaxation currents plotted as a function of voltage (*Q-V*). Individual *Q-V*s for ATB^0,+^ (blue, n = 38), GlyT2 (red, n = 53) and GlyT1 (green, n=15)). The solid lines are fit with a Boltzmann equation (2). **(D-F)** combined violin and box plot distributions of the Boltzmann equation parameters (D) *z*_max_, (E) *V*_1/2_, and (F) *Q*_max_ for GlyT1 (green, n = 15), GlyT2 (red, n = 53) and ATB^0,+^ (blue, n = 38). **Figure 3–Figure supplement 1.** Expression of ATB^0,+^ increases the linear capacitance of *Xenopus* oocytes

### ATB^0,+^ and GlyT2 share similar charge movements

We used the charge-movement of ATB^0,+^ as a proxy of their cell-surface expression in *Xenopus* oocytes. The ***Figure 3***B shows representative traces of the presteady-state currents (PSSCs) of GlyT1, GlyT2 and ATB^0,+^ rthat are ecorded in glycine-free media and isolated using their specific inhibitors (ORG24598, ORG25543 and *α*-Methyl-D,L-tryptophan (*α*MT), respectively) to subtract the oocyte endogenous-currents. It could be noticed that ATB^0,+^ and GlyT2 share asymmetric PSSCs, with large outward-currents and biphasic time-courses, whereas GlyT1-PSCCs are weakly voltage-dependent (***Figure 3***B).

The charge movement plotted as function of voltage (*Q-V*) confirms the likeness of ATB^0,+^ and GlyT2, with the same slope and clear evidence of saturation (***Figure 3***C). Fits of *Q-V*s with a Boltzmann equation (2) show similar apparent charge (*z*_max_) for ATB^0,+^ (1.26±0.02 *e*, n = 38) and GlyT2 (1.25±0.01 *e*, n = 53) that are almost one charge higher than for GlyT1 (0.37±0.02 *e*, n = 14, ***Figure 3***D). The potential of half distribution (*V*_1/2_) is positive for ATB^0,+^ (+16.2±0.9 mV, n = 38) and right-shifted relative to GlyT2 and GlyT1 (−35.0±0.8 mV (n = 5) and −17.3±3.3 mV (n = 14), respectively, ***Figure 3***E). Finally, the average ATB^0,+^-*Q*_max_ is 2.4-fold larger than for GlyT2 (44.0±2.8 nC (n = 38) *vs.* 18.6±1.3 nC (n = 53), respectively ***Figure 3***F), thus confirming ATB^0,+^ higher cell-surface expression.

Overexpression of membrane transporters can expand the cell-surface of injected oocytes (***Hirsch et al., 1996***), we compared the linear membrane capacitance (Cm) of non-injected oocytes (207±3 nF, n = 33) with oocytes expressing GlyT1 (227±7 nF, n = 11), GlyT2 (243.0±5.2 nF, n = 10), and ATB^0,+^ (301.0±6.3 nF, n = 50, p<0.001) ***Figure 3***–***Figure Supplement 1***C. As the membrane capacitance is known to be proportional to the surface area (Cm = 1 μF/cm^2^), the ~45 % increase in Cm suggests a similar expansion of surface area in oocytes expressing ATB^0,+^.

### A thermodynamic determination of ATB^0,+^ stoichiometric coefficients

To establish the stoichiometric coefficients of each cosubstrate, we adapted to ATB^0,+^ the reversal potential slope method (e.g., ***Appendix 1***) that successfully solved the stoichiometry of EAAT3, GlyT1, and GlyT2 (***Zerangue and Kavanaugh, 1996***; ***Roux and Supplisson, 2000***). For this, it was first necessary to alter the intracellular composition of oocytes by micoinjection and find a set of intracellular concentrations able to reduce the excessive driving force of ATB^0,+^ and shift its reversal potential below +50 mV.

Initial attempts with a small nanoliter-volume injection of glycine (1 M) sufficient to evoke an outward currents in GlyT1-expressing oocytes, failed to reverse ATB^0,+^ nor GlyT2 ***Figure 4***A,B. Further addition of NaCl in the pipette solution that strongly reduced ATB^0,+^ driving-force as indicated by the lower amplitude of the transport current evoked by a second glycine application, triggers small outward currents at −20 mV in GlyT2- and ATB^0,+^-expressing oocytes (***Figure 4***A,B). Eventually, we selected injection parameters that evoked robust and steady outward currents in ATB^0,+^-expressing oocytes (***Figure 4***C). These transporter-mediated currents are isolated by subtraction with *α*MT ***Figure 4***CD, and are bidirectional, with a stable and measurable reversal potential (***Figure 4***D, blue triangles)) that is sensitive to extracellular manipulation of each cosubstrate concentration.

**Figure 4.**
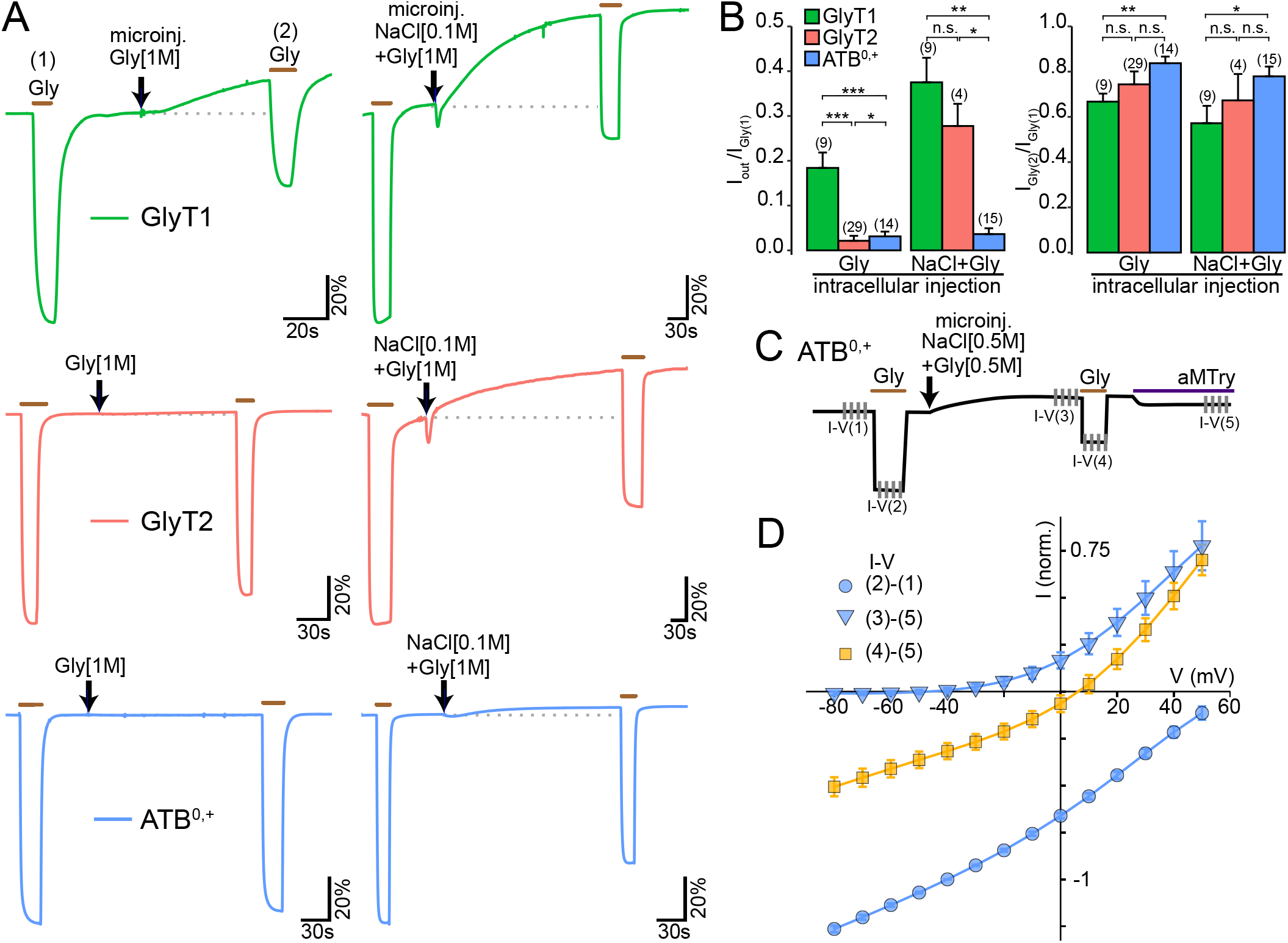
Microinjections of NaCl and glycine reverse ATB^0,+^ **(A)** representatives current traces in voltage-clamp oocytes expressing GlyT1 (top trace, green), GlyT2 (middle, red) and ATB^0,+^ (bottom, blue) and hold at V_*H*_ = −20 mV for two applications of glycine (200 μM before and after intracellular microinjection of a small volume solution(arrow: 20–40 nL). Left panels: microinjection of glycine (1 M) reverse GlyT1 as shown by the slow rise of an outward current (I_*out*_), but not GlyT2 and ATB^0,+^. Right panels: microinjection of NaCl (0.1 M) and glycine (1 M) increases I_*out*_ in GlyT1- and GlyT2-expressing oocytes but does not evoke robust I_*out*_ in ATB^0,+^-expressing oocytes, while reducing the amplitude of the current for a second glycine application. Currents are normalized by the first glycine application. **(B)** Summary data for experiments in (A). The histogram show the relative amplitude of I_*out*_ (left) and the relative current evoked by the second glycine application (right) (mean ±SEM, n are indicated in parenthesis; *** p<0.001, **p<0.01, *p<0.05; Wilcoxon rank-sum test with Bonferroni correction for multiple comparison). **(C)** Schema of the typical protocol with five I-Vs that allow recordings of outward and inward currents for ATB^0,+^ before and after microinjection of 16–20 nL of a solution containing 0.5 M NaCl and glycine. ATB^0,+^-mediated currents are isolated by subtraction of the I-V recorded in the presence of *α*-MT (500 μM). **(D)** Normalised ATB^0,+^ current-voltage relationships (IVs) for two glycine applications (200 μM) before (blue circles) and after (orange squares) 18 nL microinjection of glycine (0.5 M) and NaCl (0.5 M) that generates only outward currents (blue triangles). The IV subtractions are indicated in the legend and correspond to the five IVs described in C. Data are mean ±SEM for six oocytes. Amplitudes are normalized by the absolute amplitude of the glycine current recorded before injection at V_*H*_ =−40 mV.

Using this experimental paradigm, we impaled two electrode voltage-clamp oocytes expressing ATB^0,+^ with a third micropipette and injected 13–23 nL of solution containing 0.5 M NaCl and glycine. As a steady outward current develops following injection, we constructed three I-Vs for each oocyte with either a change in glycine (2, 0.2, and 0.02 mM, ***Figure 5***A), Na^+^ (100, 30, and 10 mM, ***Figure 5***B), or Cl^−^ (100, 30, and 10 mM, ***Figure 5***C) and the current reversal potentials were plotted in linear-log plots for each cosubstrate (***Figure 5***D). Regression analysis show similar slopes for glycine (28.6±1.0 mV/decade, n = 8) and Cl^−^ (31.5±2.7 mV/decade, n = 6), but much steeper for Na^+^ (84.7±4.5 mV/decade, n = 11 ***Figure 5***D). The average stoichiometric coefficients determined by the slope ratios (*n*_Gly_= 1.0±0.1, *n*_Cl_= 1.1±0.1, and *n*_Na_= 2.9±0.2, ***Figure 5***E) support a 3 Na^+^, 1 Cl^−^, 1 glycine stoichiometry for ATB^0,+^.

**Figure 5.**
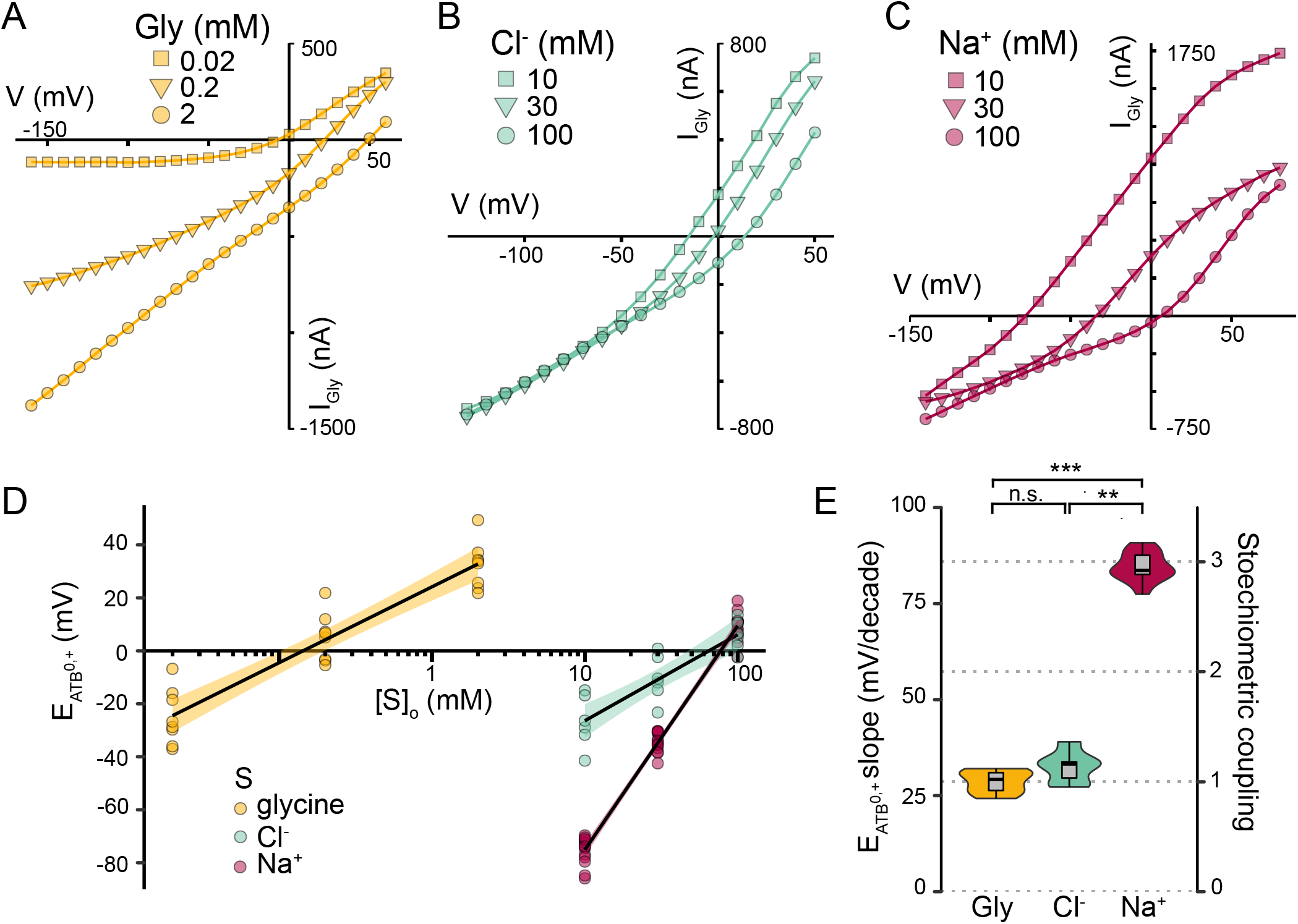
The reversal potential slopes method establishes a 3 Na^+^/1 Cl^−^/glycine stoechiometry for ATB^0,+^ **(A-C)** shifts in reversal potentials of three I-Vs recorded in microinjected ATB^0,+^-expressing oocytes (13–23 nL) of a concentrated (0.5 M) solution of NaCl + glycine) for (**(A)**) hundred-fold change in external glycine concentration or ten-fold concentration change in **(B)** Cl^−^ and **(C)** Na^+^. Currents recorded with the same protocol in the presence of *α*-MT (500 μM or 1 mM at low Na^+^) have been subtracted. **(D)** Semi-log plot of reversal potentials for the experiments described in A-C as function of each cosubstrate concentrations (S). Slopes of reversal potential changes were calculated for glycine (orange, 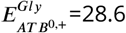, R^2^=0.873, p<2.47 10^−11^, n = 8 oocytes), Cl^−^ (green, 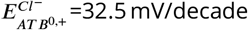, R^2^=0.754, p=2.96 10^−6^, n = 6 oocytes), and Na^+^ (red, 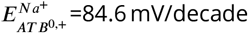, R^2^=0.983, p<2 10^−16^, n = 14 oocytes) with the 95 % confidence interval shown in shade areas. **(E)** combined violon and box plot distributions of ATB^0,+^ reversal potential slopes and stoichiometric coefficients for glycine, Cl^−^ and Na^+^. **Figure 5–Figure supplement 1.** ATB^0,+^ charge-coupling and turnover rate

In agreement, we measured a charge coupling (*z*_T_) of 2.08 *e*/glycine from the slope of the linear relationship between the time-integral of the transport current and ^14^C glycine uptake, thus confirming the tight electrogenic coupling of ATB^0,+^ transport-cycle (***Figure 5***–***Figure Supplement 1***A). Finally, an apparent turnover rate (*λ* = 18 s^−1^) was estimated from the linear relationship between *I*_max_ and *Q*_max_ (29.7 s^−1^), after correction for the ratio of glycine-coupled/glycine-uncoupled charges (see ***Figure 5***–***Figure Supplement 1***B). Together, the charge coupling and turnover rate of ATB^0,+^ well predict the current amplitude for an average expression in oocyte (*Q*_max_ = 44 nC) as 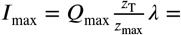 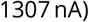, within the range reported in ***Figure 3***A.

### An external gate controls the access and locking of Na^+^ sites in the apo-ATB^0,+^

Evidence from the shift in *Q-V*s (***Figure 3***E) as well as from the abnormal rectification of the glycine-EC_50_ at negative potentials (***Figure 2***D) suggests an higher sodium affinity of the apo outward-facing conformation of ATB^0,+^, and more complex allosteric interactions with Na^+^ than for GlyT2. Therefore, we examined in more details the Na^+^- and voltage-dependence of ATB^0,+^ and GlyT2 charge-movement.

The ***Figure 6***A shows a marked and unexpected reduction in the envelope of ATB^0,+^-PSSCs for a tenfold reduction in sodium concentration. The upper range of the voltage steps was extended to +110 mV in order to catch evidence of saturation, such as the current-crossing as the charge movement approached saturation but not the current peak-amplitude (***Figure 6***A). Traces in ***Figure 6***A show that a marked current-crossing at 100 mM and 50 mM Na^+^ that is strongly reduced at 20 mM and absent at 10 mM.

**Figure 6.**
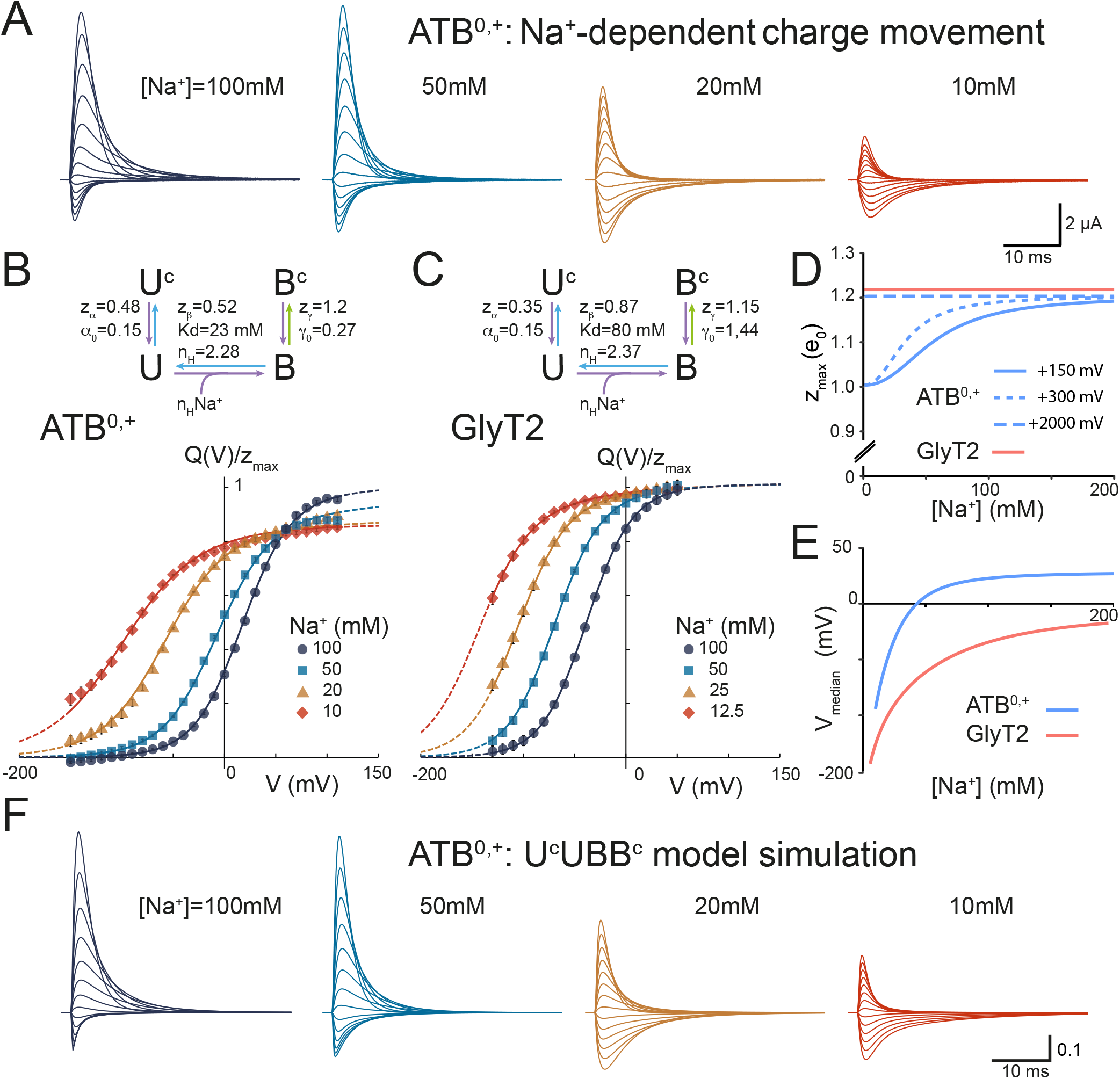
The charge movement shows evidence for an external gate controlling Na^+^ access and locking in apo-ATB^0,+^ **(A)** Representative traces of ATB^0,+^-PSSCs recorded at the onset of voltage steps (from +110 mV to −150 mV by decrements of 20 mV; V_*H*_ = −40 mV) in glycine-free solution containing the indicated Na^+^ concentration. ATB^0,+^-specific PSSCs are isolated by subtraction of traces recorded in the presence of *α*-MT (1 mM); any residual currents have been subtracted for clarity. **(B-C)** Normalyzed *Q-V*s for ATB^0,+^ (B, n = 14, ±SEM) and GlyT2 (C, n = 7, ±SEM) for the four Na^+^ concentrations shown in A, with the same color code. The data are average of sets of four *Q-V*s normalized by the *Q*_max_ value for [Na^+^]=100 mM. The solid lines correspond to the fit of the four-state scheme (#3) depicted above, with a gating, Na^+^-binding and Na^+^-locking steps that are detailed in ***Figure 6***–***Figure Supplement 1***. The green and blue arrows in the model represent the two steps activated during depolarization while the purple arrows identifies steps activated by hyperpolarizing voltage steps. The dashed lines of the *Q-V*s are extrapolations of the model outside the experimental voltage range. **(D-E)** The (D) median voltage (D) and apparent charge displacement (E) plotted as function of [Na^+^]. The median voltage is estimated as shown in (***Figure 6***–***Figure Supplement 6***). **(F)** model prediction of ATB^0,+^-PSSCs for the same conditions as in A. The rate constants are described in the Materials and Methods section. The voltage independent rate constants are k12_0_=79 s^−1^, k23_0_=88 mol^−1^ s^−1^ and k34_0_=23 s^−1^, with *d_α_* = 0.36, *d_β_* = 0.42 and *d_γ_* = 0.7. The off rate constants are back calculated using the equations, equilibrium constants and valencies detailed in ***Figure 6***–***Figure Supplement 1***B. **Figure 6–Figure supplement 1.** A four-state sequential model for the charge movement of ATB^0,+^ and GlyT2 **Figure 6–Figure supplement 2.** The equivalent charge displacement of ATB^0,+^ is Na^+^-dependent **Figure 6–Figure supplement 3.** *Q-V*s Global fits **Figure 6–Figure supplement 4.** Distribution of the four states of scheme #3 at different Na^+^ concentrations **Figure 6–Figure supplement 5.** *Q*_max_ convergence at low Na^+^ concentration required extreme voltage **Figure 6–Figure supplement 6.** Na^+^-dependency of the median voltage for *Q-V*s of ATB^0,+^ and GlyT2

According to a minimal two-state Hill model for multiple Na^+^ proposed for GAT1 (***Mager et al.*** (***1996***, 1998)), and corresponding to the scheme #1 in ***Figure 6***–***Figure Supplement 1***A), a ten-fold reduction in [Na^+^] is predicted to left-shift the *Q-V* without altering its slope nor *Q*_max_ (***Mager et al., 1996***). Although individual fits with a Boltzmann equation support this prediction for GlyT2 (***Figure 6***–***Figure Supplement 2***A-B), this is not the case for ATB^0,+^ as shown by the decreasing peaks of the *Q-V* first derivative (***Figure 6***–***Figure Supplement 2***C-D). As expected, the scheme #1 generates a poor global fit of ATB^0,+^ *Q-V*s as function of Na^+^, while being acceptable for GlyT2 ***Figure 6***–***Figure Supplement 3***A).

Then we tested a sequential, linear model that preserves Hill formalism for simplicity, with a single and voltage-dependent binding step for multiple Na^+^ with a charge *zβ*, but includes a gating step controlling the access to or priming of Na^+^-sites for binding (schemes #2 and #3, ***Figure 6***–***Figure Supplement 1***A). Gate opening and closure are voltage-dependent with *zα* the charge of the gating step (U↔U^*c*^) of Na^+^ unbound transporters and *zγ* the charge displacement that lock Na^+^ (B↔B^*c*^) (***Figure 6***–***Figure Supplement 1***A).

Because these two-path exits from the Na^+^-bound state (B) at positive potentials could carries asymmetric charges (*zγ*>*zα*+*zβ*), scheme #3 was able to solve the apparent Na^+^-dependency of *z*_max_ (***Figure 6***D) as indicated by the overwhelming difference in Akaike criterion information (ΔAICc) tabulated in ***Figure 6***–***Figure Supplement 1***B. Furthermore, the scheme #3 describes also the constant *z*_max_ of GlyT2 *Q-V* (***Figure 6***D), as *zγ ≤ zα* + *zβ*. Na^+^ locking in the apo outward facing conformation of ATB^0,+^ is further facilitated by an higher affinity for Na^+^, with a 3.5 fold difference in Kd (23 and 80 mM, for ATB^0,+^ and GlyT2, respectively), whereas conversely, GlyT2 lower affinity facilitates gate closure primarily from the Na^+^-unbound state U (***Figure 6***–***Figure Supplement 1***C,***Figure 6***–***Figure Supplement 4***). As expected for a linear model, extrapolations at extreme voltage (+2 V) confirm a convergence to the same *Q*_max_ value ***Figure 6***–***Figure Supplement 5***.

Globally, the fit parameters for ATB^0,+^ and GlyT2 in the scheme #3 show a remarkable coherency (***Figure 6***B,C), with little difference in the Hill coefficients (2.3 and 2.4, for ATB^0,+^ and GlyT2 respectively). Because it is a four-state model, the median voltage (V_*med.*_) was estimated as function of [Na^+^] (***Figure 6***D) as shown in ***Figure 6***–***Figure Supplement 6***) that is positive for [Na^+^]> 40 mM for ATB^0,+^ but always negative for GlyT2, up to 200 mM (***Figure 6***E), indicating that Na^+^ binding in the absence of voltage is energetically favorable for ATB^0,+^ while not for GlyT2 ***Figure 6***F.

Finally, we challenged the model #3 to reproduce the highly asymmetric and biphasic time courses of ATB^0,+^ transient current using the fitted equilibrium constants and charges of ***Figure 6***B (see material and method). ***Figure 6***F shows simulations that effectively recapitulate the Na^+^ dynamics of ATB^0,+^ PSSCs, supporting further the gating and locking mechanisms of scheme #3.

### 3-Na^+^ coupling limits efflux and exchange of glycine transporters

The driving force of a 3 Na^+^-coupled transporter can support large flux asymmetry that favor a net influx while limiting a net efflux, although unidirectional efflux by homo- or heteroexchange is still possible as shown for glutamate transporters (***Kavanaugh et al., 1997***). Limiting heteroexchange would be beneficial for ATB^0,+^ function considering its broad substrate specificity, by limiting futile cycles and leak of intracellular metabolites. Therefore, we compared the rate of glycine efflux and exchange in voltage-clamp oocytes pre-loaded with ^14^C-glycine and hold at V_*H*_ = −60 mV in order to minimize the basal efflux rate. The rate of efflux of ^14^C-glycine for each transporter was monitored in the outflow solution under basal condition and during glycine application (***Figure 7***A).f As expected from their difference in driving force, the basal glycine efflux was lower for ATB^0,+^ and GlyT2 compared to GlyT1 (***Figure 7***,A,B). Application of glycine stimulated exchange of GlyT1-expressing oocytes as shown by the 8-fold increase in efflux rate (***Figure 7***A,B), while only slightly in GlyT2 and ATB^0,+^-expressing oocytes (***Figure 7***A,B).

**Figure 7.**
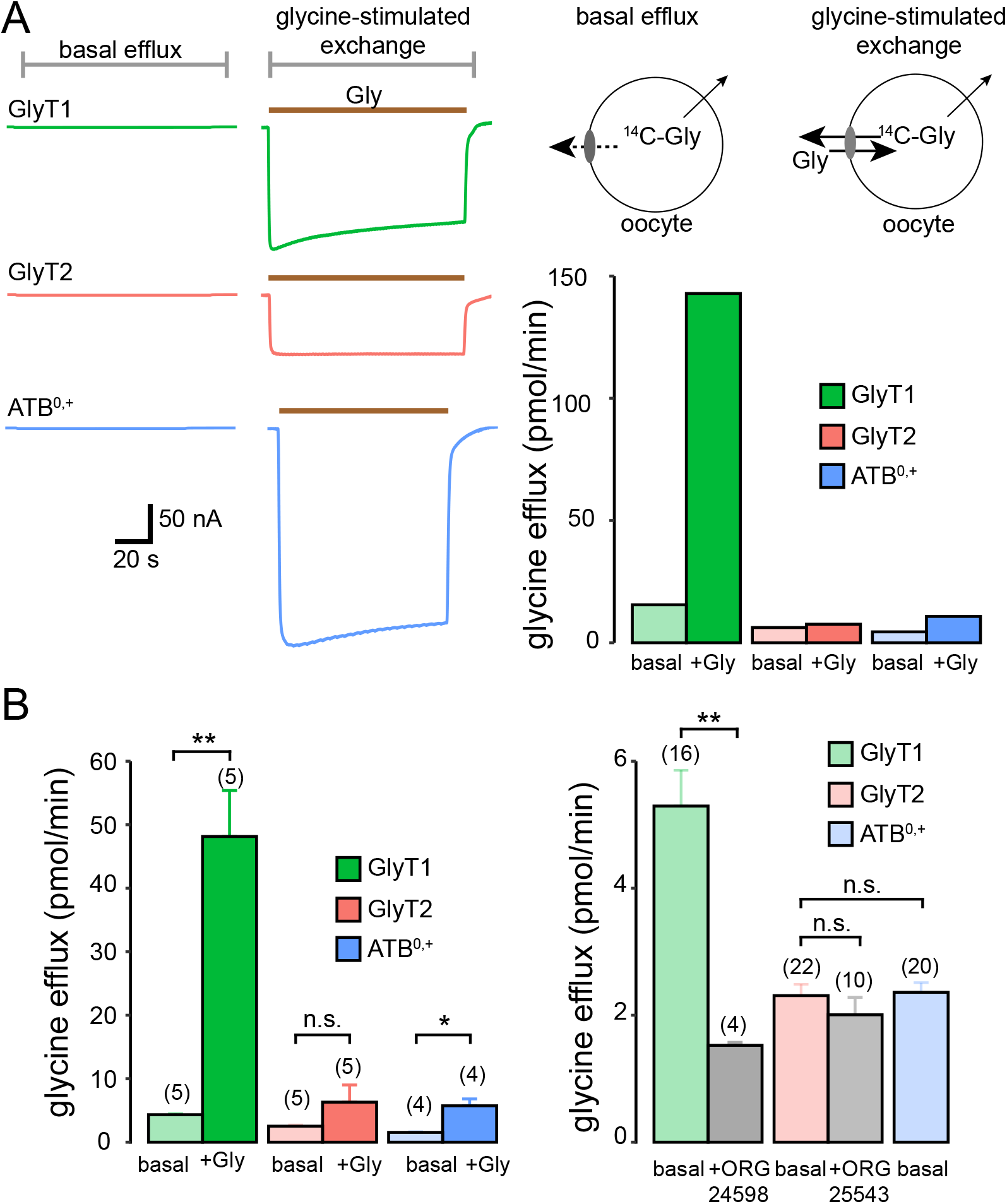
3 Na^+^-coupling limits glycine efflux and exchange **(A-C)** extracellular glycine promotes efflux in GlyT1-expressing oocytes but not in GlyT2- and ATB^0,+^-oocytes. **(A)** The efflux of glycine was measured from voltage-clamped oocytes hold at −60 mV and injected with concentrated ^14^C-glycine. The outflow of the perfusion solution was collected for one minute in the absence (basal, left traces) or in the presence of glycine (1 mM, glycine-stimulated exchange, right traces) for oocytes expressing GlyT1 (top, green traces), GlyT2 (middle, red traces) or ATB^0,+^ (bottom, blue traces). The bars histogram show glycine efflux in basal condition and during glycine application for the three transporters (same experiment). The schema shows the two components of glycine efflux in these experiments: a transporter-specific path that allow possible exchange in the presence of extracellular glycine and a non-specific component for all endogenous glycine leakage by the endogeneous transporters of *Xenopus* oocytes. **(B)** Summary data for the transtimulation of efflux (left panel) and the basal efflux (right panel). Application of OR24598 inhibits a large fraction of the basal efflux in GlyT1-expressing oocytes whereas ORG25543 has no effect, indicating minor contribution of GlyT2 to the basal glycine efflux. No inhibitor was tested with ATB^0,+^ since the basal efflux was as low as GlyT2-expressing oocytes.

## Discussion

Here, we establish a new 3 Na^+^, 1 Cl^−^, 1 amino acid stoichiometry for ATB^0,+^ that supports its unidirectional and concentrative uptake properties, and provides a rationale for its structural proximity with GlyT2 in the SLC6 family.

### A thermodynamic revision of ATB^0,+^ stoichiometry

The charge-coupling of +2.08 *e*/glycine measured in ATB^0,+^-expressing oocytes forced us to question the large consensus about its 2 Na^+^, 1 Cl^−^ stoichiometry, as this charge is larger than predicted from this stoichiometry and required either (1) a third Na^+^-coupling like GlyT2 (***Roux and Supplisson, 2000***), (2) a Cl^−^-exchange as proposed for GAT1 (***Loo et al., 2000***), or (3) a substrate-gated uncoupled-conductance as for the anion conductance of the glutamate transporters (***Wadiche et al., 1995b***; ***Cater et al., 2016***).

We used the reversal potential slope-ratio method that is based on the zero-flux equation (***Appendix 1***) and has successfully solved the stoichiometric coefficients of EAAT2-3, GlyT1-2 and GAT1 (***Zerangue and Kavanaugh, 1996***; ***Levy et al., 1998***; ***Roux and Supplisson, 2000***; ***Lu and Hilgemann, 1999***; ***Willford et al., 2015***). As previously reported for GlyT2, it was first necessary to reduce the excessive inwardly-directed driving force of ATB^0,+^ at saturating glycine concentration (2 mM), as equation 6 predicts reversal potentials of 169 mV or 111 mV for a 2 or 3 Na^+^/ 1Cl^−^-coupled transporter, respectively (assuming [Gly]_*i*_ = 1 mM, [Na^+^]_*i*_ = 10 mM and [Cl^−^]_*i*_ = 35 mM). To find a set of intracellular concentrations that allow ATB^0,+^ to operate bidirectionally, we routinely injected 22 nL solution of NaCl + glycine (0.5 M) that increases their concentration by ~17.7 mM assuming an oocyte effective water-volume of 600 nL, that is sufficient to left-shift the current reversal potential to +32.8 mV or +33.1 mV for a 2 or 3 Na^+^-coupling, respectively, which is in agreement with the average value of 32.5 mV in ***Figure 5***D.

The average reversal potential slope of 28.6 mV determined for a ten-fold reduction in [Gly]_*e*_ is close to the theoretical slope of 29.6 mV for two charges at 25 °C (Equation 6), and confirms that they are both energetically coupled to the transport cycle. The slope values for Na^+^ (84.6 mV/decade) and Cl^−^ (32 mV/decade) are also closed to the theoretical values for 3 Na^+^/gly and 1 Cl^−^/gly (88.4 and 28.6 mV/decade, respectively), thus establishing a 3 Na^+^/1Cl^−^/1 glycine stoichiometry for ATB^0,+^.

### Does ATB^0,+^ need such a large driving force ?

The increase in driving force provided by a third Na^+^ substantiates the “highly-concentrative” adjective often used to describe ATB^0,+^-mediated uptake (***Barilli et al., 2020***; ***Gupta et al., 2005***; ***Hatanaka et al., 2001***; ***Karunakaran et al., 2008***, ***2011***). Nevertheless, a weaker driving force is predicted for ATB^0,+^ than the up to a million fold glycine accumulation of GlyT2 (***Roux and Supplisson, 2000***; ***Supplisson and Roux, 2002***), because of the smaller amplitude of Na^+^ and Cl^−^ electrochemical gradients in the colon and lung epithelia than in nerve terminals.

Located in the apical membrane of colonocytes in the absorptive epithelium (***D’Argenio et al., 2006***; ***Gupta et al., 2005***; ***Hatanaka et al., 2002***; ***Ugawa et al., 2001***), ATB^0,+^ absorbs mixtures of amino acids issues from the bacterial proteolysis taking place in the outer mucus layer and lumen of the colon, and ATB^0,+^ clearance of nutrients helps to maintaining an apparent sterility of the adherent inner mucus layer (***Atuma et al., 2001***; ***Johansson et al., 2008***; ***Jakobsson et al., 2015***; ***Li et al., 2015***; ***Chen et al., 2020***).

As over 90 % of NaCl and water that enter the cecum are reabsorbed in the proximal colon (***Phillips, 1969***; ***Thiagarajah and Verkman, 2018***; ***Sandle, 1998***), considerably lower [Na^+^] (~15 mM) and [Cl^−^] (<10 mM) have been measured at the surface of the rat distal colon (***Talbot and Lytle, 2010***; ***Chen et al., 2020***). If these concentrations correspond to those of the thin mucus layer, then a 3 Na^+^-coupling is not an option but a requirement for ATB^0,+^ in order to provide enough driving force for an uphill transport of amino acids since it would be minimal or even not possible otherwise (***Chen et al., 2020***).

In the lung, ATB^0,+^ plays a similar scavenging role on the apical side of the airway epithelium (***Sloan and Mager, 1999***; ***Sloan et al., 2003***; ***Uchiyama et al., 2008***; ***Paola et al., 2017***), thus depleting essential nutrients and limiting bacteria growth in the thin and sterile airway surface layer (ASL) (***Mager and Sloan, 2003***; ***Paola et al., 2017***; ***Ruffin et al., 2020***). Short circuit currents evoked by amino acids and dependent on Na^+^ have been recorded in cultured human bronchial epithelial cells and mouse trachea epithelium (***Galietta et al., 1998***; ***Sloan et al., 2003***; ***Ahmadi et al., 2019***), although the [Na^+^] and [Cl^−^] in the thin ASL are not precisely known as two opposing models predict either a low salt condition (~50 mM) or near isotonic concentrations (***Boucher, 1999***; ***Jayaraman et al., 2001***; ***Song et al., 2003***). In any event, the driving force generated by a 3 sodium-coupling should not be a limiting factor for ATB^0,+^ scavenging function.

Upregulation of ATB^0,+^ has been detected in twelve types of solid cancers (***Sikder et al., 2017***, ***2020***) suggesting that its overexpression is the main mechanism to increase the uptake rate of glutamine or glycine, because of the dynamic *cis* and *trans* competition when ATB^0,+^ is exposed to mixture of amino acids. Furthermore ATB^0,+^ overexpression in colon cancer leads to its misrouting to the basolateral side of the epithelium with unrestrained access to the serosal pool of amino acids (***Chen et al., 2020***).

### Na^+^ binding, locking and allosteric interactions

The charge movement of ion-coupled transporters is a substrate-free signature of their membrane expression and interactions with motor ions, as shown for SGLT1 (***Parent et al., 1992***; ***Loo et al., 1993***), GAT1 (***Mager et al., 1996***; ***Bossi et al., 2002***), EAAT2 (***Wadiche et al., 1995a***), NaPi-2 (***Forster et al., 1998***), GlyT1-2 (***Roux et al., 2001***; ***Cherubino et al., 2010***), GAT3 (***Sacher et al., 2002***), and SNAT2 (***Grewer et al., 2013***). The relaxation currents evoked by positive voltage steps correspond to electrogenic steps of the partial transport cycle that are associated with Na^+^ unbinding, while conversely negative voltage steps drive Na^+^ binding/occlusion in the outward-facing conformation of the apo-transporter (***Grewer et al., 2013***; ***Hilgemann and Lu, 1999***). In addition to ion displacement, the charge movement can involve the reorientation of charged residues that are confined within the membrane electric field (***Bezanilla, 2008***, ***2018***), as demonstrated by voltage-clamp fluorometry for SGLT1 and GAT1 (***Cowgill and Chanda, 2019***; ***Li et al., 2000***; ***Loo et al., 1998***; ***Meinild et al., 2009***).

Not surprisingly, the *Q-V*s of ATB^0,+^ and GlyT2 confirms their common ionic coupling as they run parallels indicating similar charge displacement, which is almost one charge higher (+0.88 *e*) than for GlyT1, thus suggesting a net contribution of the third Na^+^. Both transporters have Hill coefficient above 2, indicating a cooperativity of the three Na^+^ sites. However, ATB^0,+^ large difference in *V*_1/2_ (+51 mV, ***Figure 3***D) and lower microscopic Kd (23 *vs.* 80 mM, ***Figure 6***B-C)) relative to GlyT2 indicate an higher affinity for Na^+^. The fraction of Na^+^-bound transporters at −40 mV is 94 % for ATB^0,+^ while only 56 % for GlyT2 (closed circles in ***Figure 6***–***Figure Supplement 4***C). We show that both the slope and *Q*_max_ of ATB^0,+^ are reduced at lower Na^+^ concentrations, thus preventing a global fit by the minimal two-state model proposed by ***Mager et al.*** (***1996***) for GAT1. Although such Na^+^-dependency is unexpecte, it is nevertheless not unique as a similar reduction in slope and *Q*_max_ was reported for GAT3, yet uninterpreted with a kinetic model (***Sacher et al., 2002***). We show that addition of a gating and locking steps with asymmetric charges solve the apparent Na^+^-dependency of ATB^0,+^ *Q-V*. A dynamic equilibrium between the two Na^+^-unbinding paths, with either its extracellular release or locking in the transporter as the external gate close, and a small imbalance between the valency of the two paths predict the reduction in *z*_max_ at low Na^+^ concentrations. Conversely, the locking step plays little role for GlyT2 because of its lower affinity favored gate closure of Na^+^-unbound transporters. It can be speculated that Na^+^-locking in ATB^0,+^, by preventing Na^+^ release, stabilize Na^+^-bound conformation at depolarized potentials.

### Structural proximity of ATB^0,+^ and GlyT2

A single serine residue (S481) in the unstructured central part of TM6, controls the substrate intolerance for N-methyl derivative of GlyT2 (***Vandenberg et al., 2007***). Indeed, sarcosine evokes a current in oocytes expressing GlyT2 mutants at this position (S481G or S481A) while introducing a serine abolishes sarcosine transport in the GlyT1 mutant G305S (***Vandenberg et al.*** (***2007***)). As expected from its intolerance to sarcosine (***Figure 1***D, ***Anderson et al.*** (***2008***)), ATB^0,+^ has a serine (S320) at the equivalent position (***Figure 1***–***Figure Supplement 2***).

Although a detailed structure of a 3-Na^+^-coupled NSS is not yet available, molecular dynamics simulations with plausible atomic models of GlyT2 based on dDAT crystal structure (***Penmatsa et al., 2013***), have identified several residues for the putative third Na^+^ binding site (***Subramanian et al., 2016***; ***Benito-Muñoz et al., 2018***). In these models, the Na1 and Na2 sites correspond to those of dDAT (***Penmatsa et al., 2013***) and LeuTaa (***Yamashita et al., 2005***), close to the substrate site, allowing a direct coordination with the carboxyl of glycine, as found for leucine in the high-resolution structure of LeuTaa (***Yamashita et al., 2005***). In contrast, the proposed Na3 site is located more distantly from the substrate site and involves an electrostatic interaction with the glutamate E650 in TM10 (***Subramanian et al., 2016***; ***Benito-Muñoz et al., 2018***). It is worth noticing that ATB^0,+^ shares the same glutamate residue at this position (E489, ***Figure 1***–***Figure Supplement 2***), whereas GlyT1 has a methionine (M470) that cannot engage in an electrostatic interaction with Na^+^. The other putative residues proposed to coordinate the third Na^+^ (M278, W265, A483 (***Subramanian et al., 2016***) or E250 (***Benito-Muñoz et al., 2018***)) are all identical for the three glycine transporters, suggesting that E489 is the primary specific candidate for the Na3 site of these two glycine transporters.

## Methods and Materials

### Molecular biology

#### Heterologous expression of SLC6 transporters in *Xenopus* oocytes

The cDNAs encoding the mouse ATB^0,+^ (gift of V. Ganapathy, ***Hatanaka et al.*** (***2001***)), the rat GlyT2a (gift of B. Lopez Corcuera and C. Aragon, ***Liu et al.*** (***1993***)) and the rat GlyT1b (gift of K Smith, Synaptic, ***Smith et al.*** (***1992***)) were subcloned in a modified pRC/CMV vector as described previously in ***Supplisson and Bergman*** (***1997***). Capped cRNAs were synthesized using the Ambion mMessage mMachine T7 transcription kit (Thermo Fisher Scientific) and kept at 1 μg/μL at −80 °C.

Females *Xenopus lævis* were anesthetized using 0.1 % Tricaine methanesulfonate (MS222, SIGMA) disolved in 0.1 % NaHCO_3_ and maintained on ice. Defolliculated oocytes were harvested from ovaries by shaking incubation in Ca^2+^-free OR-2 solution (in mM: 85 NaCl, 1 MgCl_2_, 5 Hepes, pH 7.6 with KOH) containing 5 mg mL^−1^ (type A, ROCHE). Oocytes were kept at 19 °C in a Barth’s solution (in mM: 88 NaCl, 1 KCl, 0.41 CaCl_2_, 0.82 MgSO_4_, 2.5 NaHCO_3_, 0.33 CaNO_3_, 5 Hepes, pH 7.4 (with NaOH)) containing 50 μg/mL gentamycin. cRNAs (50 ng) were injected using a nanoliter injector (World Precision Instruments).

#### Sequences alignment and phylogenetic tree

The phylogeny of SLC6 transporters in ***Figure 1***A includes 63 vertebrates orthologs of *SLC6A7*, *SLC6A9*, *SLC6A5* and *SLC6A14* from cartilaginous fishes (*Callorhinchus milii* [9, 5, 14], *Rhincodon typus* [9], bony fishes (*Danio rerio* [9, 5, 14], *Oncorhynchus mykiss* [9, 14], *Lepisosteus oculatus* [7], *Hippocampus comes* [9, 5], *Nothobranchius furzeri* [9, 5, 14]), *Latimeria chalumnae* [5], amphibians (*Nanorana parkeri* [7, 5, 14], *Xenopus lævis* [9, 5], *Xenopus tropicalis* [7, 14]), reptiles (*Chelonia mydas* [7, 9, 5, 14], *Gekko japonicus* [7, 9, 5, 14], *Notechis scutatus* [7, 5, 14]), birds (*Serinus canaria* [7, 9, 5, 14], *Empidonax traillii* [7, 9, 14], *Apteryx rowi* [7, 5, 14], *Gallus gallus* [7, 9, 14]), mammals (*Monodelphis domestica* [7, 9, 5, 14], *Mus musculus* [1, 2, 3, 4, 5, 6, 7, 8, 9, 11, 12, 13, 14], *Macaca macaca* [7, 9, 14], *Macaca menestrina* [5], *Homo sapiens* [7, 9, 5, 14]) and was generated by PhyML on phylogeny.fr (https://www.phylogeny.fr/, (***Dereeper et al., 2008***)). The tree was created and color coded using iTol (https://itol.embl.de, (***Letunic and Bork, 2007***)). The sequence alignment of ***Figure 1***–***Figure Supplement 2*** was produced using PROMALS3D (http://prodata.swmed.edu/promals3d/promals3d.php). Area-proportional Venn diagrams were drawn with eulerAPE software (http://www.eulerdiagrams.org/eulerAPE/, ***Michalec et al.*** (***2014***)).

### Electrophysiology

We used O725C amplifier (Warner Instrument) for two-electrode voltage-clamp recordings in *Xenopus* oocytes. Glass electrodes filled with 3 M KCl solution had a typical resistance of 1 MQ. Experiments were performed at room temperature with oocytes perfused continuously with a solution containing (in mM): 100 NaCl, 1.8 CaCl_2_, 1 MgCl_2_, 5 Hepes, pH 7.2 (with KOH). For intracellular microinjections, oocytes were impaled with a third micropipette connected to a nanoliter injector (World Precision Instruments) and injected with small volumes (range: 13–69 nL) of concentrated glycine and NaCl. Stationary currents were digitally filtered at 10 Hz and acquired at 100 Hz using a 1320 digidata (Molecular devices). Presteady-state currents were evoked by short voltage steps 200–400 ms in the range 110–−150 mV by decrements of 10 mV. Currents were typically filtered at 1–2 kHz and digitized at 2.5–10 kHz, except for a subset of experiments where the first millisecond of the relaxation currents were precisely isolated by recording the full range of the oocyte capacitive currents at 20–50 kHz sampling frequency and filter accordingly at 4–10 kHz). Transporter currents were isolated by subtraction of traces recorded in the presence of their specific inhibitors (GlyT1: 20 μM OR24598; GlyT2: 10 μM ORG25543; ATB^0,+^: 0.5–1 mM *α*-MT).

#### Ionic coupling

Oocytes expressing ATB^0,+^ were voltage-clamped at −40 mV and perfused with a solution containing 30 μM glycine [U-^14^C] (specific activity 3.81 MBq μmol^−1^, Amersham Pharmacia Biotech) and 20 μM unlabeled glycine. After 30–120 s application and complete wash-out, unclamped oocytes were washed 3 times with ice-cold solution and lysed in 0.5 mL triton (2 %). The radioactivity was counted with a Kontron Betamatic and the charge coupling (*z_T_*) was calculated from the slope of the relationship between glycine uptake (U_Gly_) and the time integral of the glycine evoked current (Q_Gly_) (***Roux and Supplisson, 2000***).

### Chemicals

Chemicals were obtained from SIGMA Aldrich. ORG25543 and 0RG24598 were generous gift from Organon. We used choline^+^ or Tris^+^ (2-amino-2-hydroxymethyl-1,3-propanediaol) and gluconate^−^ or methyl-sulfonate^−^ as substitutes for Na^+^ and Cl^−^, respectively.

### Fit procedures

The slope and saturation of the *Q-V*s depend on the apparent charge *z*_max_ expressed in elementary charge (*e*), and the maximal charge *Q*_max_ expressed in nC is proportional to the number of transporters (Tn) :

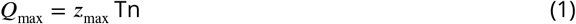

Transporter *Q-V*s in ***Figure 3*** are fitted with a Boltzmann equation (2) to extract *Q*_max_, *z*_max_ and the half-distribution potential *V*_1/2_ (***Bezanilla and Villalba-Galea, 2013***):

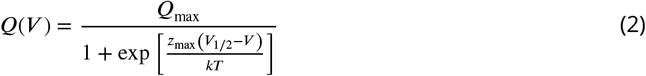

k is the Boltzmann constant, and T the absolute temperature.

In the ***Figure 6***C, sets of four *Q-V*s were measured at different Na^+^ concentrations for each oocyte, and then normalized by *Q*_max_ at [Na^+^] =100 mM using equation (2). The data set for ATB^0,+^ consists of n = 1512 samples (14 oocytes × 4 *Q-V*@[Na^+^] /oocyte × 27 voltages/*Q-V*), and n=513 samples for GlyT2 (7 oocytes × 4 *Q-V*@[Na^+^] /oocyte × 19 voltages/*Q-V*; one *Q-V*@Na^+^ = 10 mM was excluded because of a subtraction artifact). Concentrations of Na^+^ were not identical for both transporters for no particular purpose. A global fit of each data set as function of Na^+^ and voltage was performed using the equilibrium equations of schemes #1, #2, and #3 that are detailed in ***Figure 6***–***Figure Supplement 1***B, using the NonLinearFitModel function in Mathematica 12 (Wolfram). The LlAkaike values tabulated in ***Figure 6***–***Figure Supplement 1***B are the Akaike values given by Mathematica after subtraction of the minimum value for the scheme #1. The RSquared of model #3 were 0.991 and 0.990 for ATB^0,+^ and GlyT2, respectively.

In the scheme #3, *z*_max_ is not constant for ATB^0,+^ because *z_γ_ > z_α_* + *z_β_*. We computed *z*_max_ as function of Na^+^ in ***Figure 6***D, using the probability (***Zifarelli et al., 2012***) of Na^+^-unbound states (*p*(*U^c^*), *p*(*U*)) and Na^+^-bound states (*p*(*B*), *p*(*B^c^*)) as shown in ***Figure 6***–***Figure Supplement 4***:

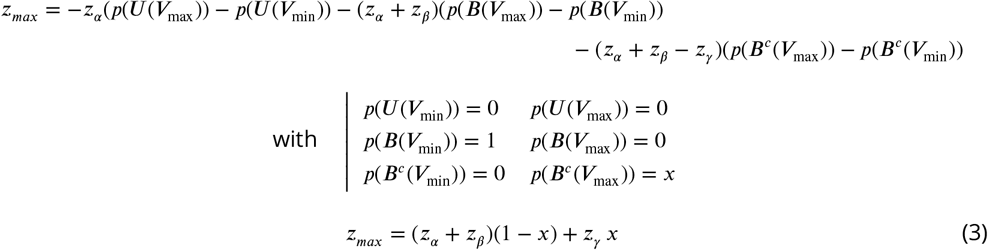

The median voltage for activation (*V*_med._) corresponds to the voltage for equal negative and positive integrals of the *Q-V* (***Figure 6***–***Figure Supplement 5***) :

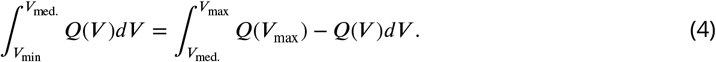

*V*_med._ is estimated using the FindRoot and Nintegrate functions of Mathematica 12.

To simulate the time course of the relaxation currents in ***Figure 6***F, we used the NDSolve function (Mathematica 12) and numerically integrates the three differential equations that defined the dynamics of the four-state model U^*c*^ UBB^*c*^ (***Figure 6***–***Figure Supplement 1***A):

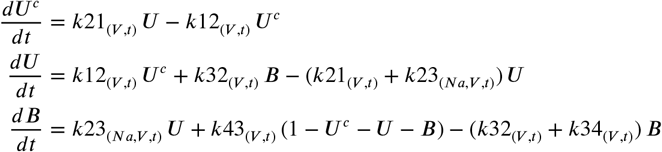

The individual rates are defined by the voltage-independent rates (*k*12_0_, *k*23_0_, *k*34_0_) and constants (*α*_0_, Kd, *γ*_0_, see ***Figure 6***–***Figure Supplement 1***A) obtained from the *Q-V* fit (***Figure 6***B). The equivalent charge for each step (*z_α_*, *z_β_*, *z_γ_*) are expressed in *e*, and the fraction of the membrane potential (*d_α_*, *d_β_*), *d_γ_*)) define the voltage dependency of forward and bacward reactions:

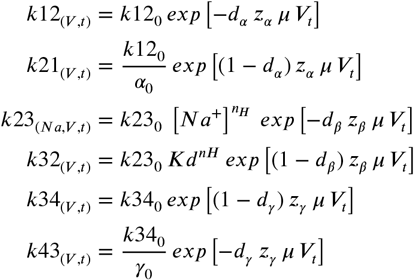

with 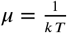, and an exponential rise of the voltage step with a time constant *τ_c_* = 300 *μ*s

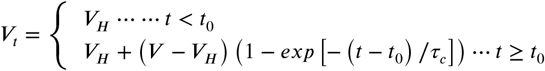

Simulations were filtered by one-order Bessel filter with cutoff frequency of 1.1154 kHz using the BesselFilterModel function of Mathematica.

## Acknowledgments

The authors wish to thanks Vadivel Ganapathy for the generous gift of the mATB cDNA. Anne Le Goff, Laurine Bonet and Annick Ayon for technical help in molecular biology. Alain Péchard and Yvon Cabirou for building the oocyte chambers and the microinjection setup. We wish to thank Sebastien Santini - CNRS/AMU IGS UMR7256 for managing the Phylogeny.fr site. We thank Stéphane Dieudonné for stimulating discussions and generous support.

## Additional information

### Author contributions

Bastien Le Guellec, Investigation, Methodology, Data Curation, Formal Analysis; France Rousseau Investigation, Data Curation, Formal Analysis; Marion Bied, Investigation, Data Curation, Formal Analysis; Stéphane Supplisson, Conceptualization, Data Curation, Formal Analysis, Investigation, Methodology, Resources, Software, Supervision, Validation, Visualization, Writing-original draft, Writing-review & editing

### Ethics

Animal experimentation: All procedures involving experimental animals were performed in accordance with the European directive 2010/63/EU on the Protection of Animals used for Scientific Purposes, the guidelines of the French Agriculture and Forestry Ministry for handling animals, and local ethics committee guidelines.

## Appendix 1 Thermodynamics of ATB^0,+^

The thermodynamic equation for ATB^0,+^ transport cycle is given by the sum of the electro-chemical gradients for all cotransported substrates:

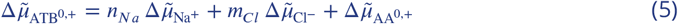

where *n_Na_* amd *n_Cl_* are the stoichiometric factors for Na^+^ and Cl^−^, respectively and AA^0,+^ designed any amino acid substrate of ATB^0,+^. In the case of a neutral susbtrate acid such as glycine, eq (5) simplify to:

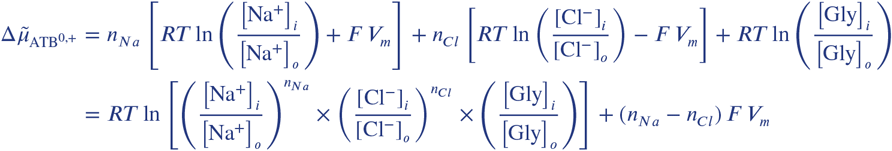

where R is the universal gas constant (8.314 J mol^−1^), T is the absolute temperature (298 °K) and F is the Faraday’s constant (96 485 C mol^−1^).

The reversal potential of ATB^0,+^ 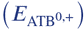 is given by:

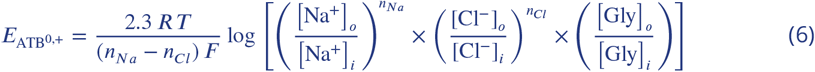

The slopes of the log-linear relationships for each cosubstrate is the product of a constant 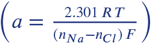 and the stoichiometric coefficients:

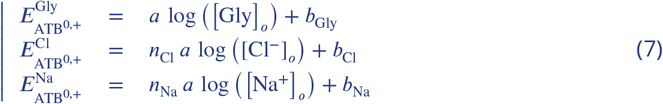

---

**Figure 1–Figure supplement 1.**
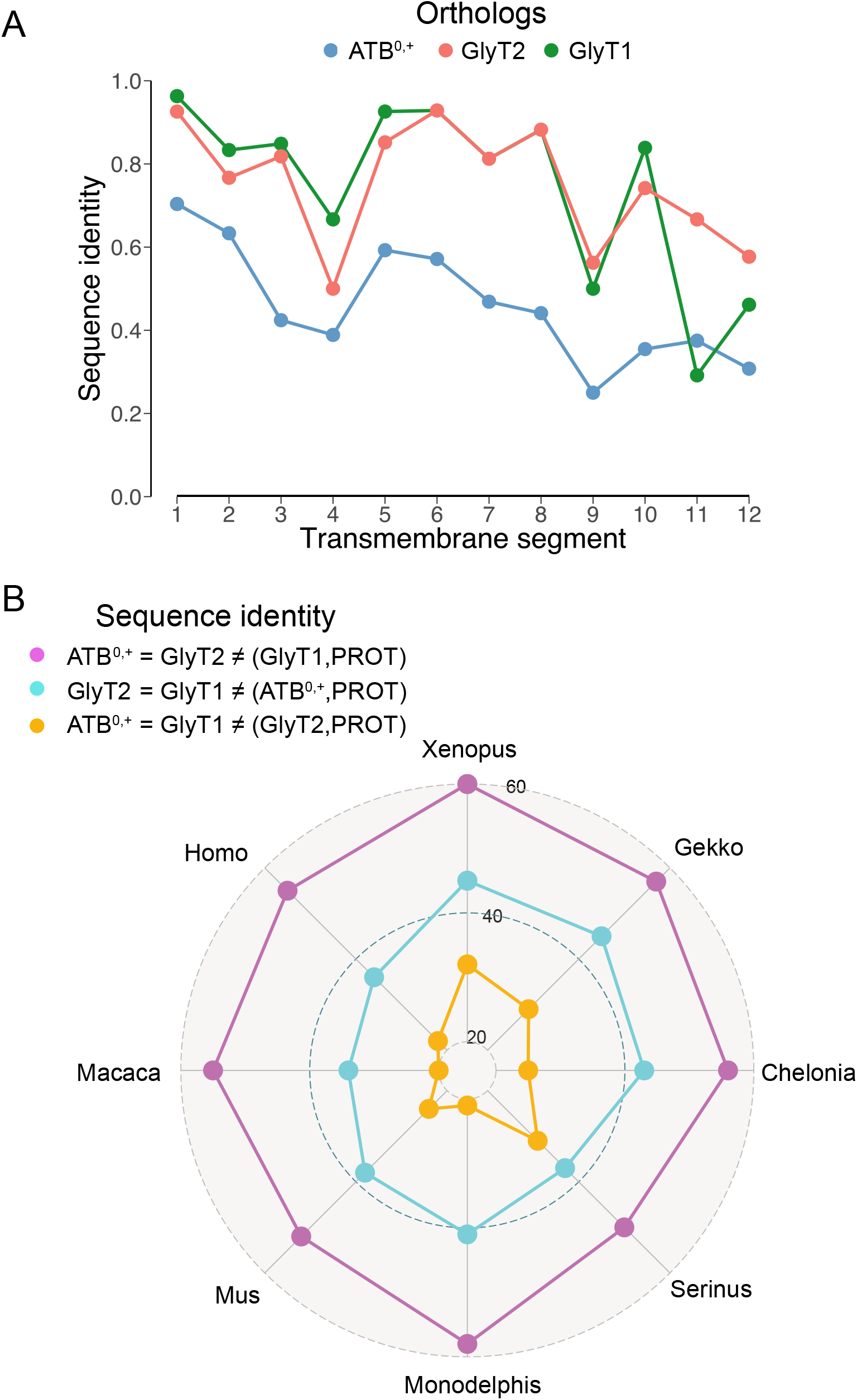
**(A)** Fraction of sequence identities in the transmembrane segments (TMs) for all Vertebrates orthologs shown in **Figure 1**A, using the human SERT structure as a template for TMs delimitation(***Coleman et al., 2016***). **(B)** Radar plot of the cumulative number of sequence identity for each glycine-transporter pairs (ATB^0,+^-GlyT2 (purple), GlyT2-GlyT1 (cyan), ATB^0,+^-GlyT1 (yelow)). Pair identity shared with PROT are excluded as indicated in the legend

**Figure 1–Figure supplement 2.**
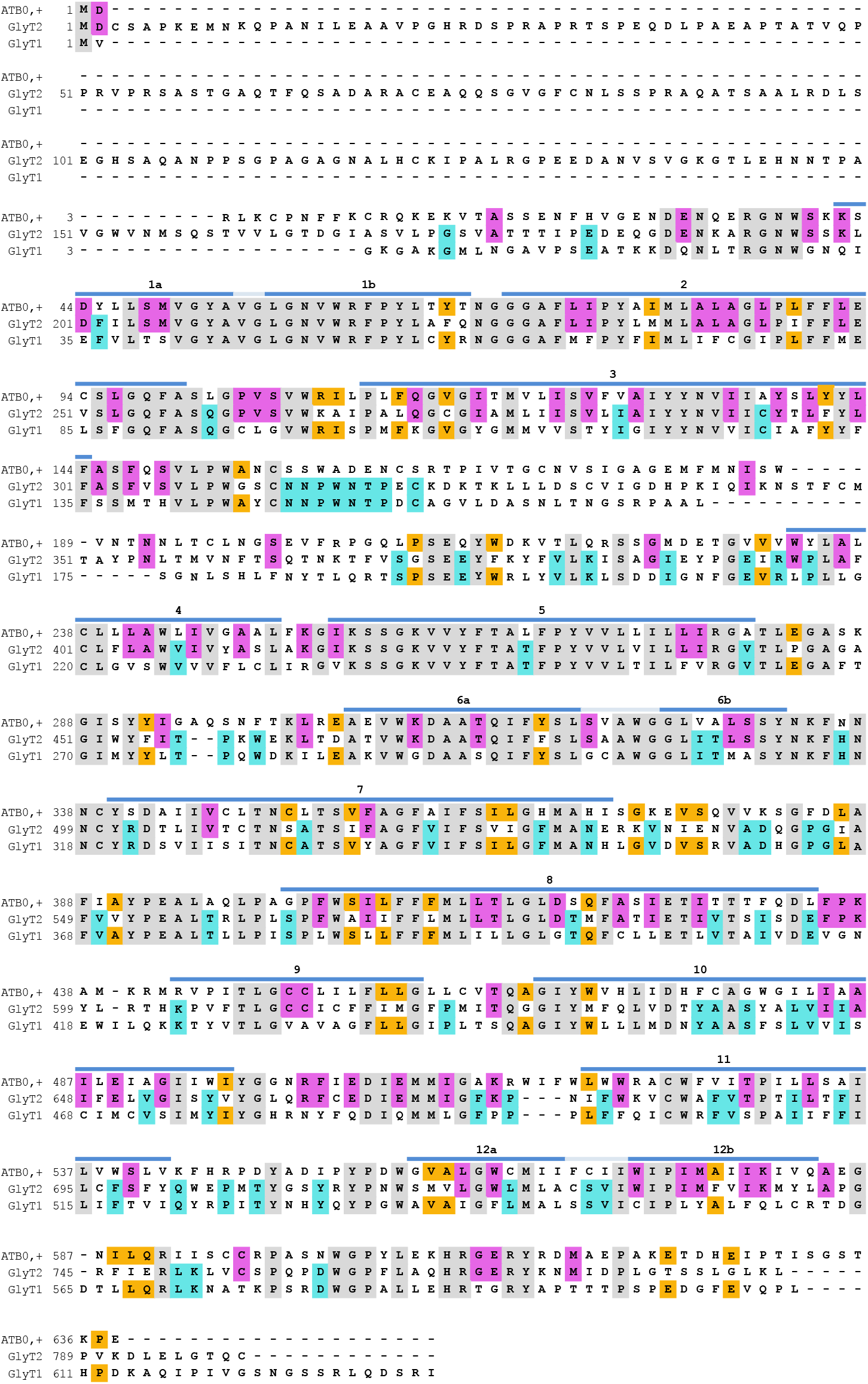
Sequences alignment of the mATB^0,+^, mGlyT2 and mGlyT1. The color code indicates amino acids common for the three transporters (gray, n = 220), and the identity shared between ATB^0,+^ and GlyT2 (purple, n = 95), ATB^0,+^ and GlyT1 (cyan, n = 43), GlyT1 and GlyT2 (orange, n = 76).

**Figure 2–Figure supplement 1.**
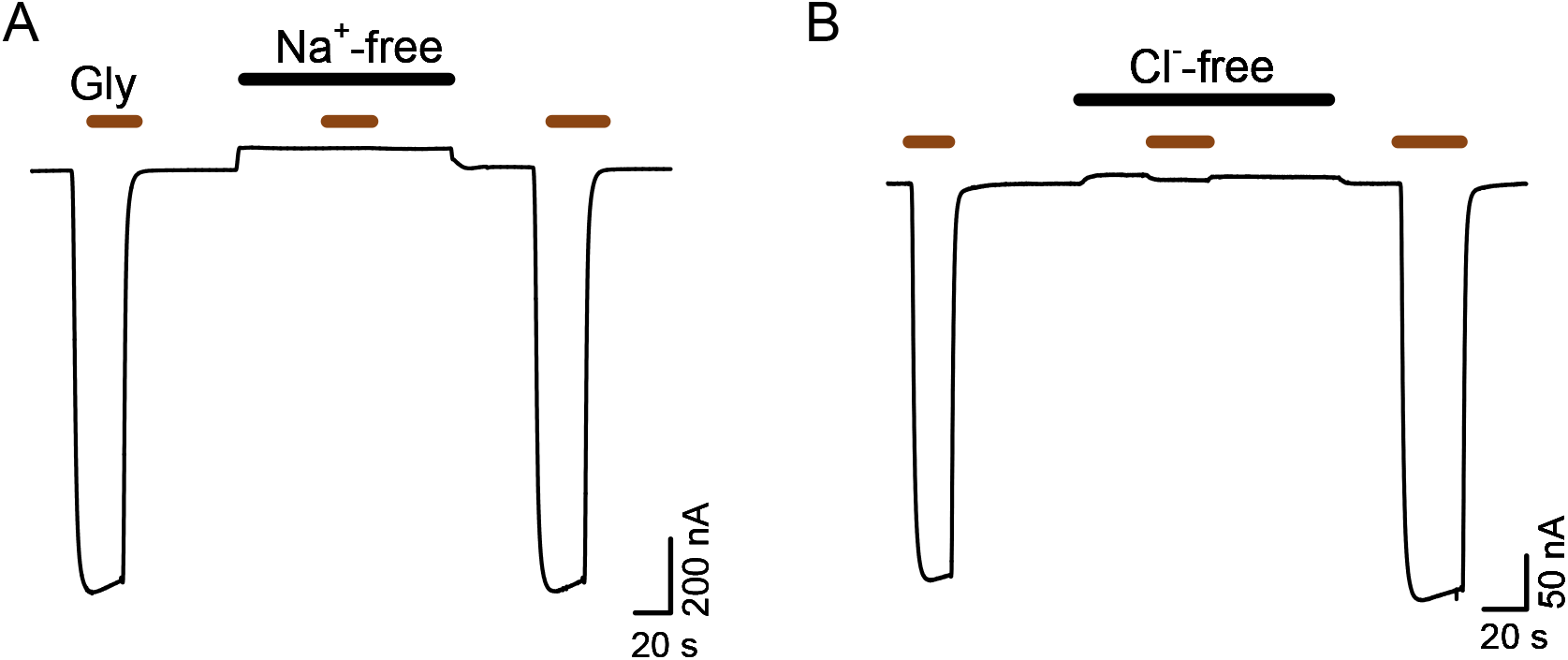
Glycine evoked current in ATB^0,+^-expressing oocytes is Na^+^- and Cl^−^- dependent (red bar: [Gly]=200 μM, V*_H_* = −40 mV). Na^+^ was substituted by choline (left trace, solid bar) and Cl^−^ was substituted by gluconate (right trace, solid bar).

**Figure 3–Figure supplement 1.**
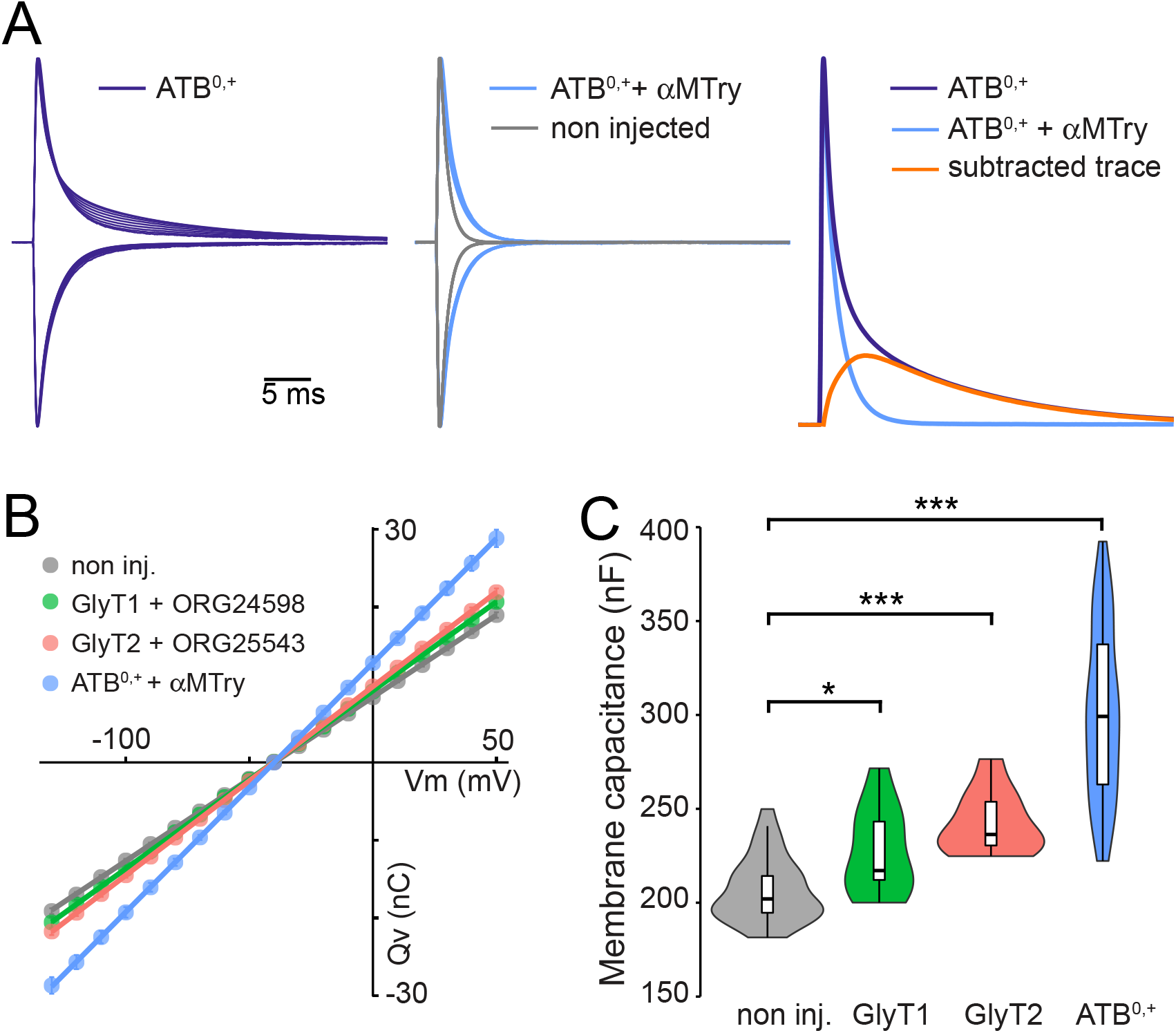
**(A)** Left panel: traces of capacitive currents (linear and PSSCs) recorded at the onset of voltage steps (from +50 mV and −130 mV by decrements of 10 mV) and normalized by the peak amplitude for a representative ATB^0,+^-expressing oocyte. Each trace is normalized by the absolutepeak current (the trace at V*_H_* = −40 mV is not included). Middle panel: same as left in the presence of 1 mM *α*-MT (blue traces) to block ATB^0,+^-charge movement. The gray traces represent currents recorded with the same protocol for a representative non injected oocyte. Right panel: the charge movement of ATB^0,+^ (orange trace) for a single step at 50 mV after *α*-MT subtraction. **(B)** Cm is given by the slope of the linear charge-voltage relationships and increases in oocytes expressing transporter membrane capacitance (Cm) of non-injected oocytes (209±3 nF, n = 33) with oocytes expressing GlyT1+ORG24598 (227±7 nF, n = 11), GlyT2 +ORG25543 (242.8±5.2 nF, n = 10), and ATB^0,+^ + *a*-MT320.6±11.7 nF, n = 16). **(C)** Combined violin and box plots of the distribution of linear membrane capacitance of non-injected oocytes (n = 33), and oocytes expressing GlyT1 (n = 11), GlyT2 (n = 10) and ATB^0,+^ (n = 50; a group of 34 ATB^0,+^-expressing oocytes was added by estimating the slope of the *Q-V* at hyperpolarized potentials).

**Figure 5–Figure supplement 1.**
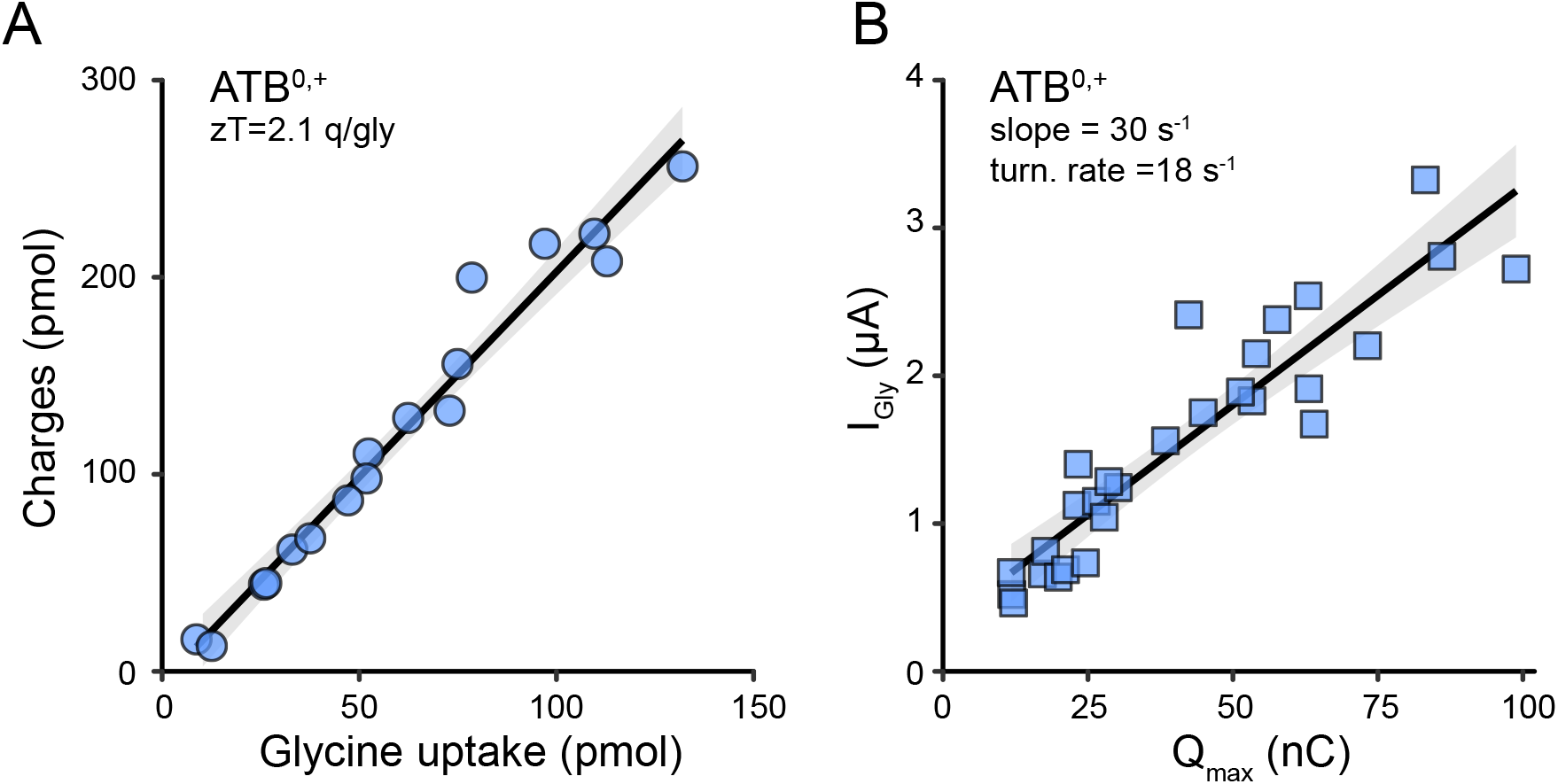
**(A)** linear relationship between glycine uptake and the time integral of the glycine-evoked current for 30-120 s application of ^14^C-glycine (50 μM, each circle represents a single oocyte expressing ATB^0,+^). The slope (*z_T_* =2.08 *e*, R^2^=0.982, p=2.16 10^−12^) gives a charge coupling per glycine in agreement with the 3 Na^+^/1 Cl^−^ stoichiometry. The shaded area shows the 95 % confidence interval. **(B)** linear relationship between Imax ([Gly] = 1–2 mM) and *Q*_max_ for oocytes expressing ATB^0,+^ and hold at V*_H_* =−40 mV. An apparent turnover rate (*λ*=18 s^−1^) is derived from the *I*_max_/*Q*_max_ slope (29.7 s^−1^, R^2^=0.848, p=3.77 10^−12^) using the equation 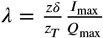, with z_*T*_ =2.08 *e* and z*δ*=1.26 *e* as determined in A and **Figure 3**D). The shade area corresponds to the 95 % confidence interval.

**Figure 6–Figure supplement 1.**
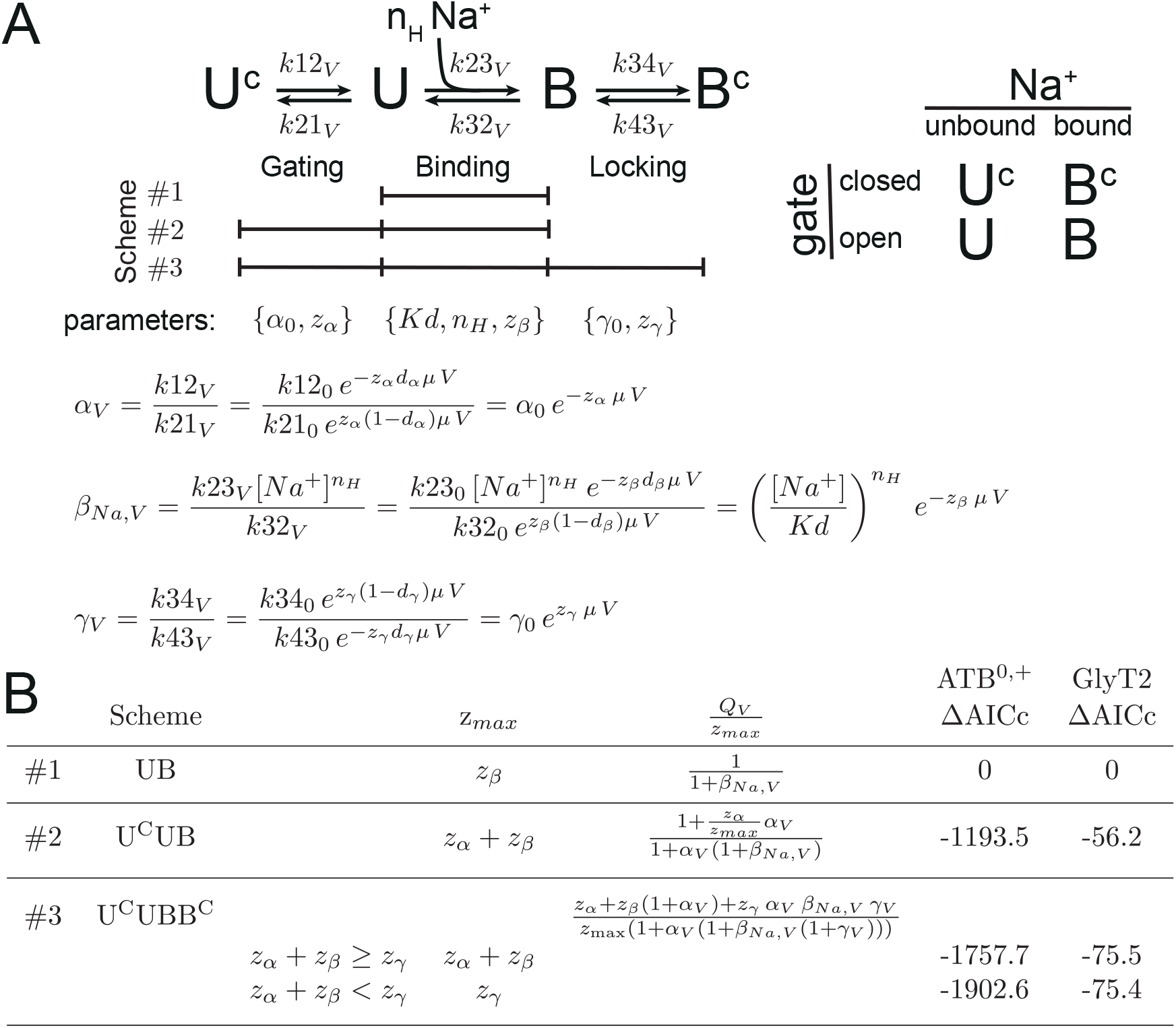
**(A)** The model shows the three kinetic schemes tested here to fit ATB^0,+^- and GlyT2-*Q-V*s as function of Na^+^. In glycine-free media, outward facing transporters are assumed to be either Na^+^-unbound (U^*c*^, U) or Na^+^-bound (B, B^*c*^). The UB scheme (#1) corresponds to the binary Hill model proposed by ***Mager et al.*** (***1996***) for GAT1 and is limited to a single binding step for multiple Na^+^. It is defined by 3 parameters: the microscopic dissociation constant Kd, the Hill coeffcient *n_H_*, and the equivalent charge *z_β_*. For simplicity, we define *β_Na,V_* as the equilibrium constant of the binding step as function of Na^+^ and voltage. Schemes #2 and #3 include a gating step controlling the access of Na^+^ to its binding sites. The equilibrium constants are *α_V_* and *γ_V_* for the gating and locking steps, respectively. In the scheme #2, gate closure occurs only in the unbound state (U) whereas it is independent of Na^+^-binding in scheme #3. At negative potentials all transporters are in B state (***Figure 6***–***Figure Supplement 4***). Positive voltage steps trigger gate closure from Na^+^-unbound (U) and Na^+^-bound state (B). **(B)** The *z*_max_ and normalized *Q-V* equations are tabulated for each scheme, with the Δ*AICc* values estimated from the fit. *z*_max_ for scheme #3 is conditioned by equality or inequality of the valencies of the Na^+^-release and Na^+^-locking paths

**Figure 6–Figure supplement 2.**
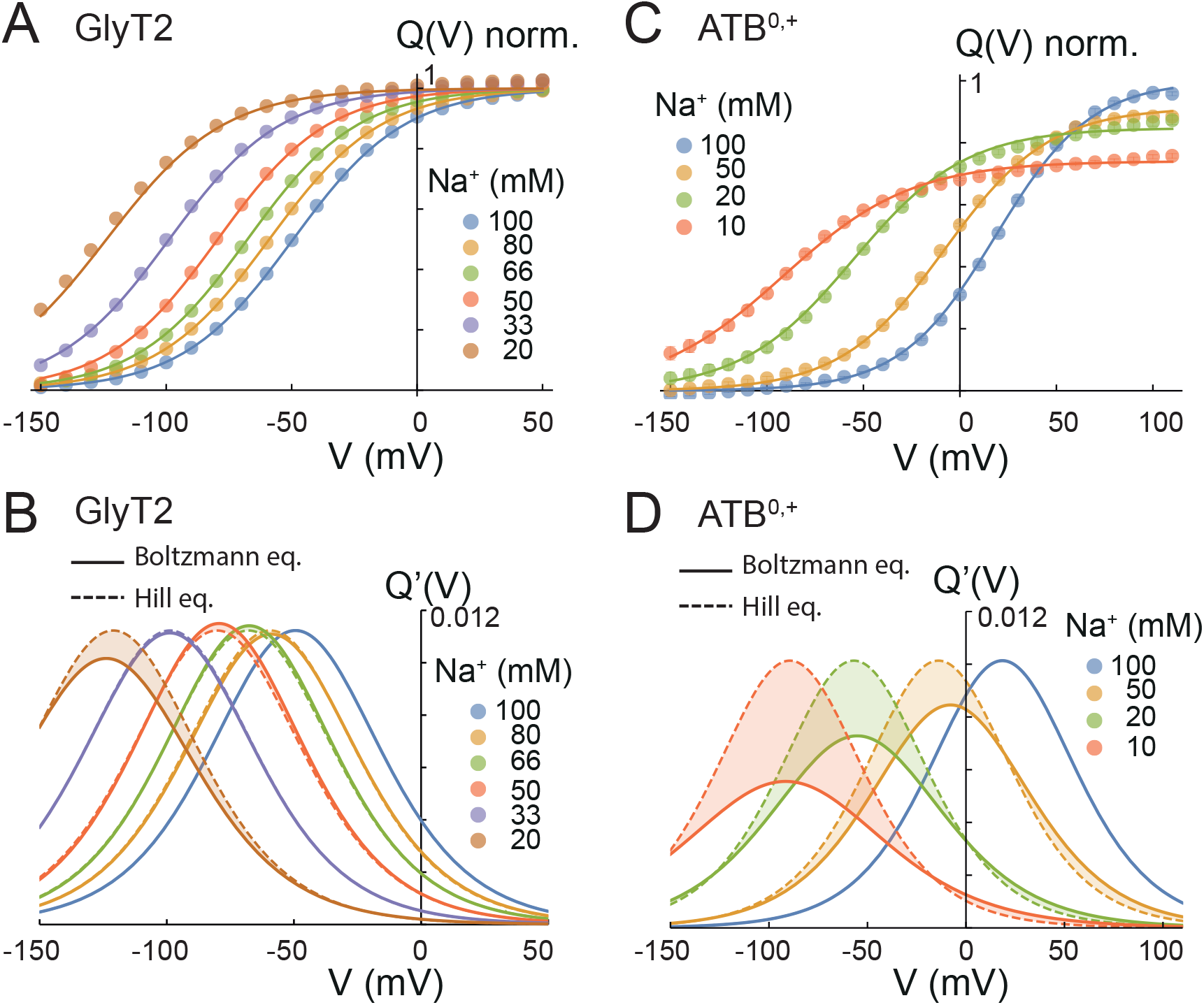
**(A-B)** *Q-V*s for ATB^0,+^ (A, n = 7) and GlyT2 (B, n = 1) for different Na^+^ concentrations indicated in the legend. The solid line corresponds to the individual fit using a Boltzmann equation (**Figure 3**C). All *Q-V*s are normalized by the *Q*_max_ value determined for Na^+^ = 100 mM . **(C-D)** The solid lines represent plots of the first derivative of the Boltzmann fits shown in A-B for ATB^0,+^ (C) and GlyT2 (D). The dashed lines are the first derivative of Hill equation (scheme #1 in ***Figure 6***–***Figure Supplement 1***) with the parameters n= 1.9, *z*_max_ =0.548 and KNa= 67.4 mM for ATB^0,+^ and n_H_ = 2.04, *z*_max_ =1.135 and Kd=290 mM for GlyT2

**Figure 6–Figure supplement 3.**
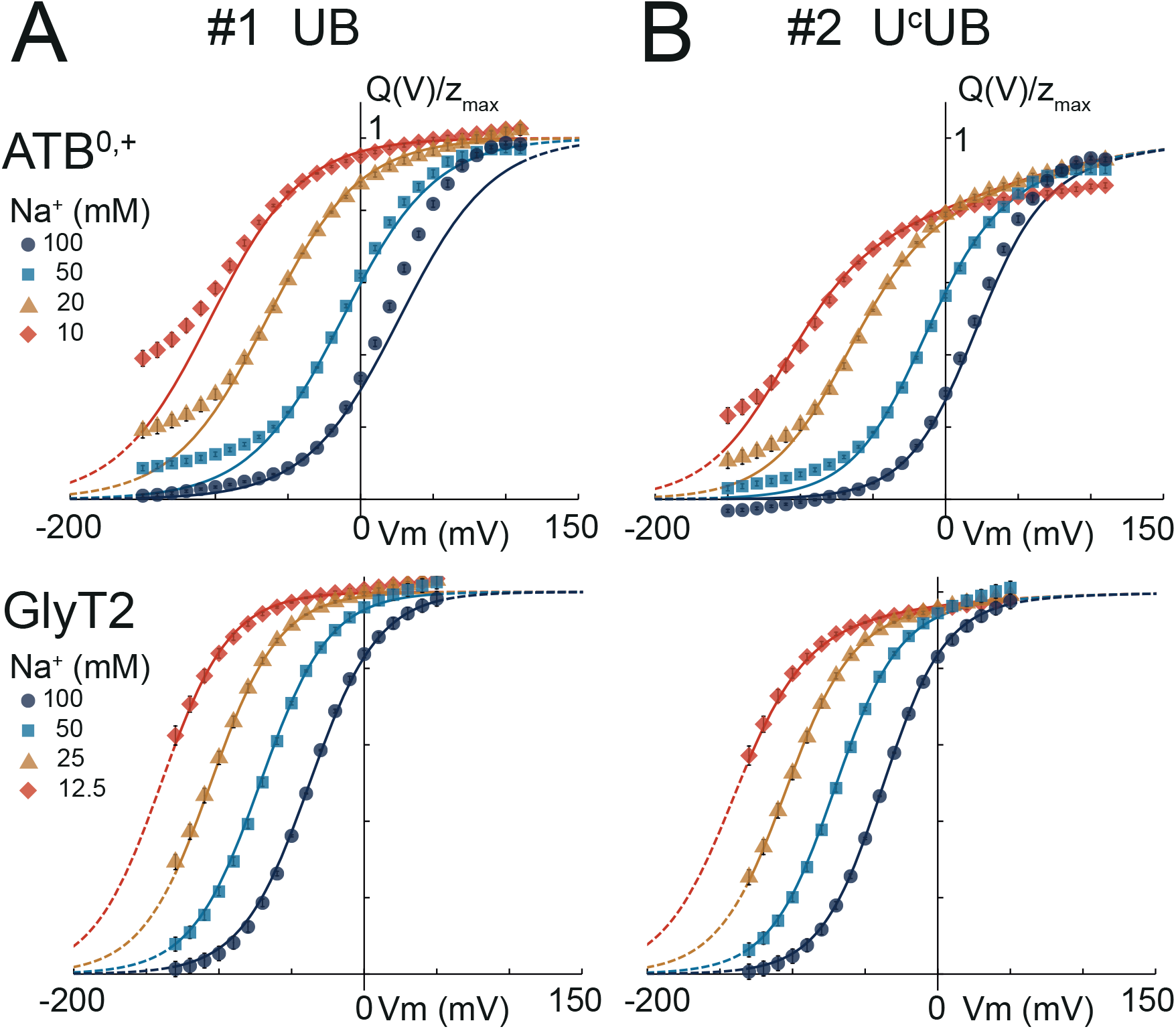
**(A-D)** Global fits of the *Q-V*s of ATB^0,+^ (top) and GlyT2 (bottom) as a function of the Na^+^ concentration for the kinetic schemes #1 (A), #2 (B) as described in ***Figure 6***–***Figure Supplement 1***.

**Figure 6–Figure supplement 4.**
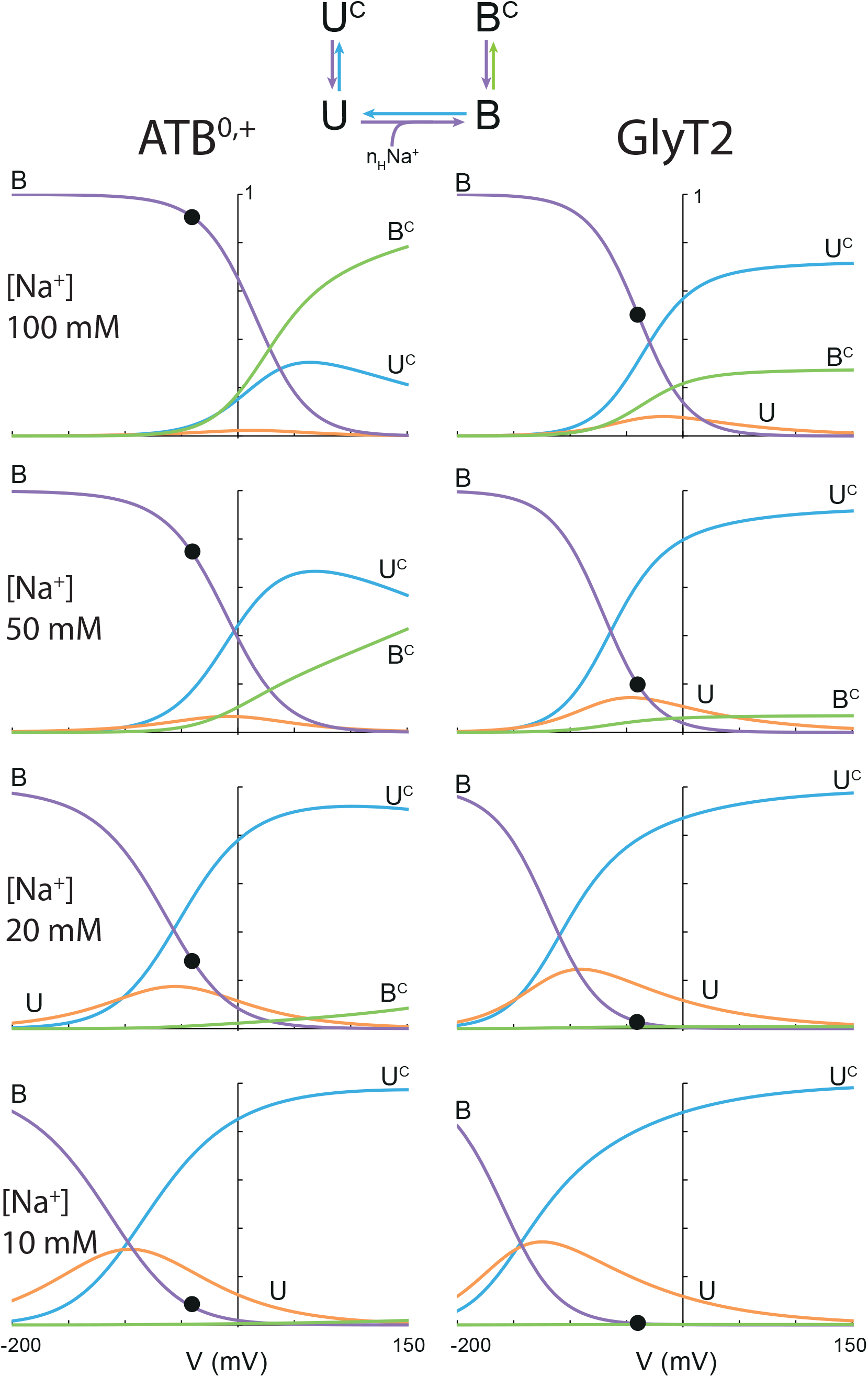
The distribution of the four-state (U^*c*^, U, B, B^*c*^ ) of ATB^0,+^ (left panels) and GlyT2 (right panels) plotted as a function of voltage for the Na^+^ concentration indicated on the left, using the set of parameters in **Figure 6**B-C. The closed circles represent the fraction of the B state at V_*H*_ = −40 mV.

**Figure 6–Figure supplement 5.**
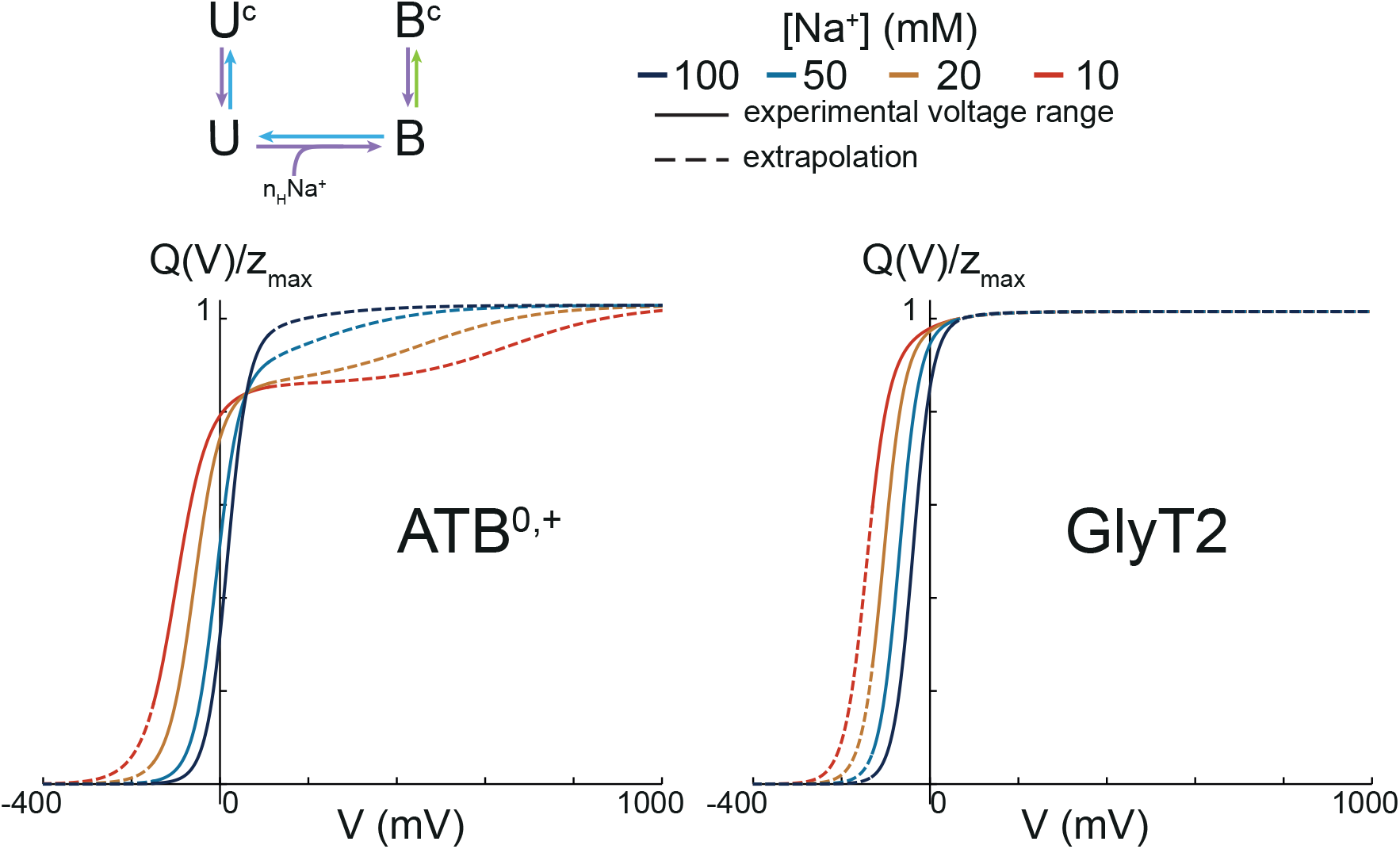
Prediction from the scheme #3 shows a reduction of the apparent *Q*_max_ at low Na^+^ concentration **Figure 6**D. Here, we show the extrapolation of ATB^0,+^- and GlyT2 *Q-V*(s) calculated for an extreme voltage range (−0.4–1 V)using the same set of parameters depicted in **Figure 6**B-C. As expected for the equilibrium equation of scheme #3, all four *Q-V*s converge to 1 at extreme positive potential.

**Figure 6–Figure supplement 6.**
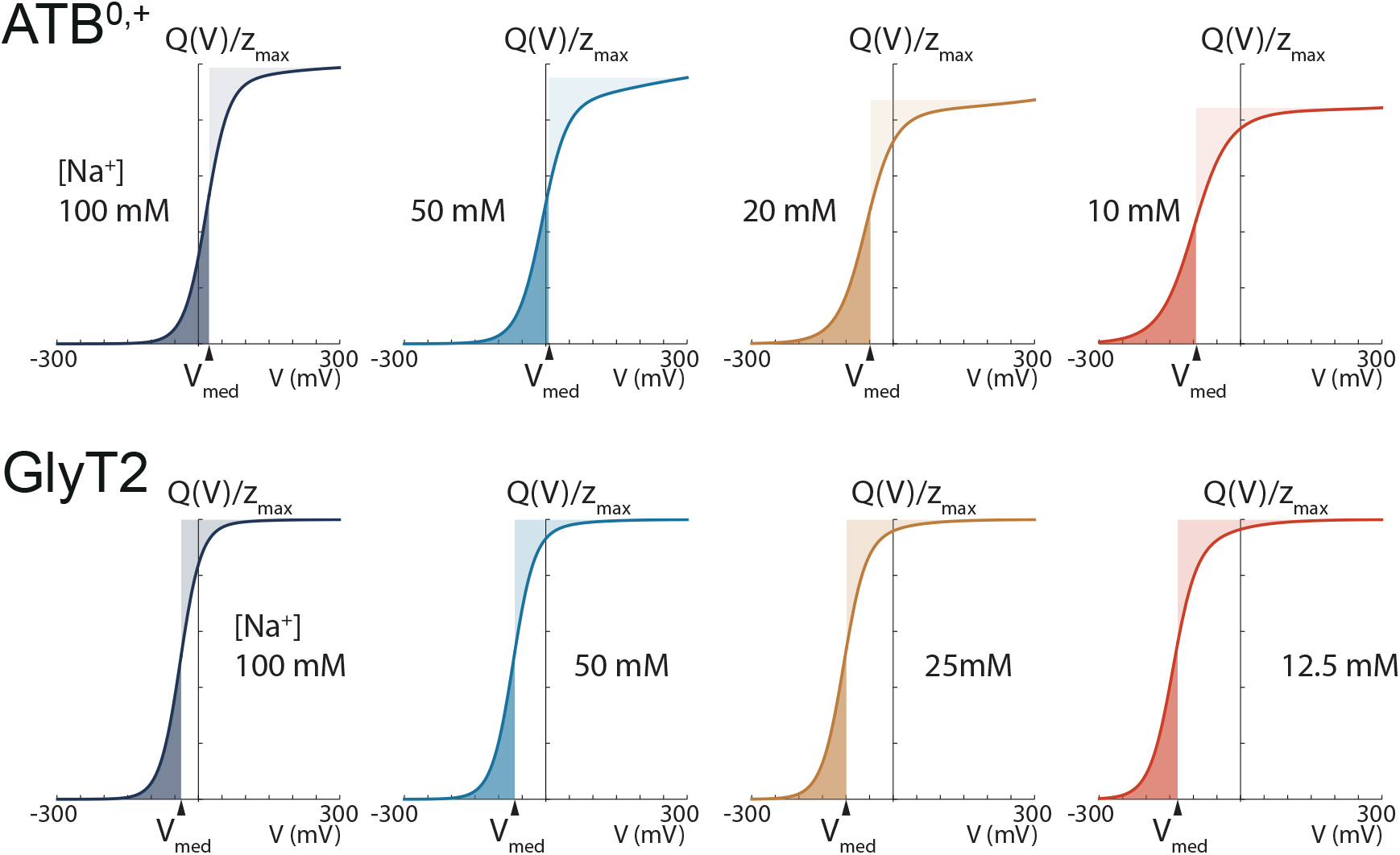
The *Q-V*s of ATB^0,+^ (top) and GlyT2 (bottom) are plotted for four Na^+^ concentrations as indicated, using the fit parameters of scheme #3. The median voltage (V_*med.*_) for equal charge distribution is estimated by numerical integration as described in methods and indicated on the voltage axis of each plot. The darker and lighter shade areas represent the negative and positive integrals, respectively.

